# SIMBA: SIngle-cell eMBedding Along with features

**DOI:** 10.1101/2021.10.17.464750

**Authors:** Huidong Chen, Jayoung Ryu, Michael E. Vinyard, Adam Lerer, Luca Pinello

## Abstract

Recent advances in single-cell omics technologies enable the individual and joint profiling of cellular measurements. Currently, most single-cell analysis pipelines are cluster-centric and cannot explicitly model the interactions between different feature types. In addition, single-cell methods are generally designed for a particular task as distinct single-cell problems are formulated differently. To address these current shortcomings, we present *SIMBA*, a graph embedding method that jointly embeds single cells and their defining features, such as genes, chromatin accessible regions, and transcription factor binding sequences into a common latent space. By leveraging the co-embedding of cells and features, SIMBA allows for the study of cellular heterogeneity, clustering-free marker discovery, gene regulation inference, batch effect removal, and omics data integration. SIMBA has been extensively applied to scRNA-seq, scATAC-seq, and dual-omics data. We show that SIMBA provides a single framework that allows diverse single-cell analysis problems to be formulated in a unified way and thus simplifies the development of new analyses and integration of other single-cell modalities. SIMBA is implemented as an efficient, comprehensive, and extensible Python library (https://simba-bio.readthedocs.io) for the analysis of single-cell omics data using graph embedding.

## Introduction

Technology to profile single cells has advanced to several molecular modalities, dramatically advancing our ability to characterize cell states as well as discover key molecular machinery that underlies both development and disease. Individual cells are now measured using multiple molecular modalities, simultaneously. At the same time, single-cell experiments have scaled such that tens of thousands of cells can be routinely profiled. The emergence of single-cell multi-omics technologies allows for the measurements of multiple cellular layers, including genomics, epi-genomics, transcriptomics, and proteomics. Such assays have pioneered an avenue toward a better understanding of the interplay between layers as they jointly define cell states based on diverse genomic and molecular features including genes, regulatory elements, and transcription factors. While single-cell multi-omic assays have quickly evolved towards the incorporation of additional modalities with increasing resolution, harnessing their full potential has posed several significant computational challenges.

Many single-cell computational methods have been developed for the analysis of one modality (e.g., scRNA-seq or scATAC-seq analysis) ^1–4^. Common to these methods is a workflow that includes routine steps such as feature selection, dimension reduction, clustering, and differential feature detection. These “cluster-centric” analysis methods rely on accurately defined clustering solutions to discover meaningful and informative marker features. Unfortunately, clustering solutions may range widely within the space of the user-defined clustering resolution (number of clusters) and the chosen clustering algorithm. These parameters may markedly influence the resulting cluster assignment and clusters may not always correspond to the correct or intended cell populations, thereby leading to inconsistent and potentially misleading biological annotations^5^. Although initial efforts have been made recently to develop clustering-free approaches to discover informative genes, they are specifically designed for extracting gene signatures ^6, 7^ or identifying perturbations between experimental conditions^8^ from scRNA-seq data, and are therefore limited to single-modality and single-task analysis.

In addition to single-batch/modality analysis, approaches have also been proposed for multi-batch and cross-modality analysis, such as multimodal analysis (distinct cellular parameters are measured in the same cell)^9^, batch correction (the same cellular parameter is measure in different batches) ^10–12^, and integration of multi-omics datasets (distinct cellular parameters are measured in different cells)^11, 12^. These approaches play a critical role in removing batch effects that confound true biological variation, improving the characterization of cell states by leveraging the unique strengths of each assay, and providing insights into the complex mechanisms of gene regulation. However, these tasks are formulated differently from those in single-batch/modality settings and thus require development of new dedicated analysis techniques. Also, while multiple types of cellular features might be present, the relation between features cannot be exploited directly by most current methods. Furthermore, similar to single-batch/modality analysis methods, these methods identify marker features based on groups of cells obtained by clustering and therefore are limited to clustering solutions. Additionally, instead of directly identifying marker features in the integrated space, most batch correction/multi-omics integration methods need to first detect marker features in the uncorrected/unintegrated original space of each batch/modality independently, and then combine them, thus resulting in potentially inconsistent interpretations between batches/modalities.

To overcome the limitations in both single-batch/modality analysis and multi-batch/cross-modality analysis, we propose SIMBA (**SIngle-cell eMBedding Along with features**), a versatile single-cell embedding method that co-embeds cells and features into a shared latent space, in which various types of tasks can be performed based on the proximity between entities including cells and features such as genes, peaks, and DNA sequences. Unlike existing methods that require featurization of cells, SIMBA directly encodes the cell-feature or feature-feature relations into a large multi-entity graph. For each task, SIMBA constructs a graph, wherein differing entities (i.e., cells and features) are represented as nodes and relations between these entities are encoded as edges. Once the graph is constructed, SIMBA then applies a multi-entity graph embedding algorithm derived from social networking technologies as well as a Softmax-based transformation to embed the nodes/entities of the graph into a common low-dimensional space wherein cells and features can be analyzed based on their distance. Hence SIMBA provides an information-rich embedding space containing cells and all the features, serving as an informative database of entities. Depending on the task, we can define biological queries on the “SIMBA database” by considering neighboring entities of either a cell (or cells) or a feature (or features) at the individual-cell and individual-feature level (**Methods**). For example, the query for a cell’s neighboring features can be used to identify marker features (e.g., marker genes or peaks) or to study the interaction between features (e.g., peak-gene) while the query for features’ neighboring cells can be used to annotate cells.

By formulating single-cell analyses as multi-entity graph embedding problems, we show SIMBA can be used to solve popular single-cell tasks in a unified framework that would otherwise require the development of distinct specialized approaches for each task, including: 1) dimensionality reduction techniques for studying cellular states; 2) clustering-free marker detection based on the similarity between single cells and features; 3) Single-cell multimodal analysis and the study of gene regulation; 4) batch correction and omics integration analysis as well as the simultaneous identification of marker features. SIMBA is adapted to these diverse analysis tasks by simply modifying how the input graph is constructed from the relevant single-cell data.

We extensively tested SIMBA in multiple scRNA-seq, scATAC-seq and dual-omics datasets covering popular single-cell tasks including scRNA-seq analysis, scATAC-seq analysis, multimodal analysis, batch correction, and multi-omics integration. We demonstrate that SIMBA learns the joint low-dimensional representations of both cells and features and thus enables the ability to simultaneously study cellular heterogeneity as well as proximity-based marker feature detection or gene regulation inference in a clustering-free way. We also demonstrate that SIMBA performs better than or comparably to current state-of-the-art methods specifically developed for each task.

Importantly, we developed a scalable and comprehensive Python package that enables seamless interaction between graph construction, training with PyTorch for graph embedding, and post-training analysis. The SIMBA package not only provides a self-contained framework that covers preprocessing, graph embedding, and visualization, but also is compatible with popular single cell analysis tool Scanpy^2^. SIMBA with detailed documentation and extensive tutorials is available at https://simba-bio.readthedocs.io.

We believe that SIMBA, as a broadly applicable approach for single cell omics study, not only outperforms current cluster-centric analysis, but also will simplify the burden of developing methods for new single-cell tasks and measurements, while increasing interpretability of cellular mechanisms and functions.

## Results

### Overview of SIMBA

SIMBA is a single-cell embedding method with support for single- or multi-modality analyses that embeds cells and their associated genomic features into a shared latent space, generating interpretable and comparable embeddings of cells and features. It leverages recent graph embedding techniques that have been successful in modeling complex and hierarchical information present in natural languages, social networks, and other domains, as “knowledge graphs”. In our case, the graph encodes cells, different components of cellular regulatory circuits, and the relations between them.

SIMBA first encodes different types of entities such as cells, genes, open chromatin regions (peaks or bins), transcription factor (TF) motifs, and *k*-mers (short sequences of a specific length, *k*), into a single graph (**Fig. 1**, **Methods**) where each node represents an individual entity and edges indicate relations between entities. For example, if a gene is expressed in a cell, an edge is created between the gene and cell. The weight of this edge is determined by the gene expression level. Similarly, an edge is added between a cell and a chromatin region if the region is open in this cell, or between a chromatin region and a TF motif if the TF motif is found in the region.

**Figure1.**
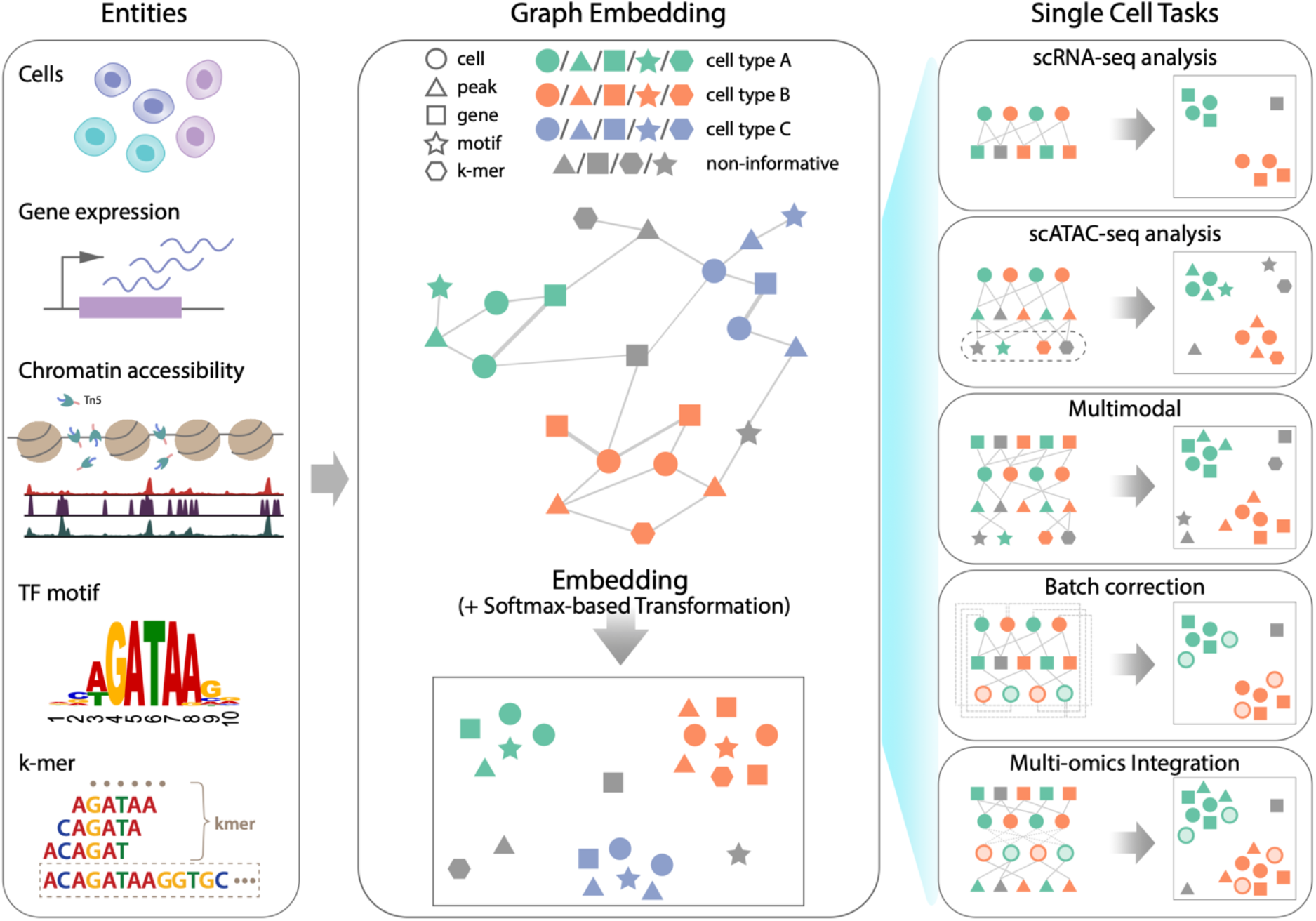
SIMBA framework overview. SIMBA co-embeds cells and various features measured during single-cell experiments into a shared latent space to accomplish both common tasks involved in single-cell data analysis as well as tasks, which remain as open problems in single-cell genomics. **(Left)** Examples of possible biological entities may be encoded by SIMBA including cells, gene expression measurements, chromatin accessible regions, TF motifs, and k-mer sequences found in reads. **(Middle)** SIMBA embedding plot with multiple types of entities into a low-dimensional space. All entities represented as shapes (cell = circle, peak = triangle, gene = square, TF motif = star, k-mer = hexagon) are colored by relevant cell type (green, orange, and blue in this example). Non-informative features are colored dark grey. Within the graph, each entity is a node, and an edge indicates a relation between entities (e.g., a gene is expressed in a cell, a chromatin region is accessible in a cell, or a TF motif/k-mer is present within an open chromatin region, etc.). Once connected in a graph, these entities may be embedded into a shared low-dimensional space, with cell-type specific entities embedded in the same neighborhood and non-informative features embedded elsewhere. **(Right)** Common single-cell analysis tasks that may be accomplished using SIMBA.

Once the input graph is constructed, a low-dimensional representation of the graph nodes is then computed using an unsupervised graph embedding method. This graph embedding procedure leverages the PyTorch-BigGraph framework ^13^, which allows SIMBA to scale to millions of cells (**Methods**). The obtained SIMBA space provides an intuitive way to study gene regulation and the regulatory mechanisms underlying cell differentiation and specification. The resulting joint embedding of cells and features not only reconstructs the heterogeneity of cells but also allows for the discovery of the defining features for each individual cell without relying on a clustering solution, separating cell-type specific features from the non-informative features. In fact, the relationship between cells and features can be explored directly through their mutual proximity in the SIMBA embedding as the distance between embedded nodes reflects their edge probability, which is informative of the potential importance of a feature to a cell and the interplay between features (**Methods**).

Therefore, cell-type-specific features such as marker genes, cis-regulatory elements can be discovered without clustering in two different ways. When the labels of cells are known, marker features can be identified as the neighboring features of cells by performing biological queries (**Methods**). When these labels are unknown, marker features can be identified through calculating the imbalance of edge probabilities between a feature and all cells using metrics such as the Gini index (**Methods**).

Importantly, graph construction is inherently flexible, enabling SIMBA to be applied to a wide variety of single-cell tasks. In the following sections, we demonstrate the application of SIMBA to several popular single-cell tasks including scRNA-seq, scATAC-seq, multimodal analysis, batch correction and multi-omics integration (**Fig. 1**). Extensions to additional tasks will become readily apparent to the reader and are later discussed.

### SIMBA enables simultaneous learning of cellular heterogeneity and individual-cell-level marker genes in scRNA-seq analysis

Single-cell RNA sequencing (scRNA-seq) is the most robust and widely used measurement to profile single cells. **Figure 2a** provides an illustrative overview of the SIMBA graph construction and the resulting low-dimensional embedding matrix of both cells and genes. Here we show how SIMBA enables simultaneous dimensionality reduction and clustering-free marker gene detection in scRNA-seq analysis. We applied SIMBA to a popular PBMCs dataset from 10x Genomics (**Supplementary Table 1**) to illustrate its workflow. After the standard preprocessing steps including normalization and log-transformation, SIMBA discretizes the gene expression matrix into multiple gene expression levels (five levels, by default). The input graph is then constructed wherein two types of nodes –cells and genes are connected by edges that embody the relation between them and are weighted according to the corresponding multiple levels of gene expression. SIMBA then generates embeddings of these nodes through a graph embedding procedure (**Fig. 2a; Methods**). Depending on the task, we have the full flexibility to visualize either the whole SIMBA embeddings (embeddings of cells and all genes in **Supplementary Fig. 1c**) or the partial SIMBA embeddings (embeddings of cells in **Fig. 2b,** or embeddings of cells and variable genes in **Fig. 2c**, or embeddings of any entities of interest) using visualization tools such as UMAP.

**Figure 2.**
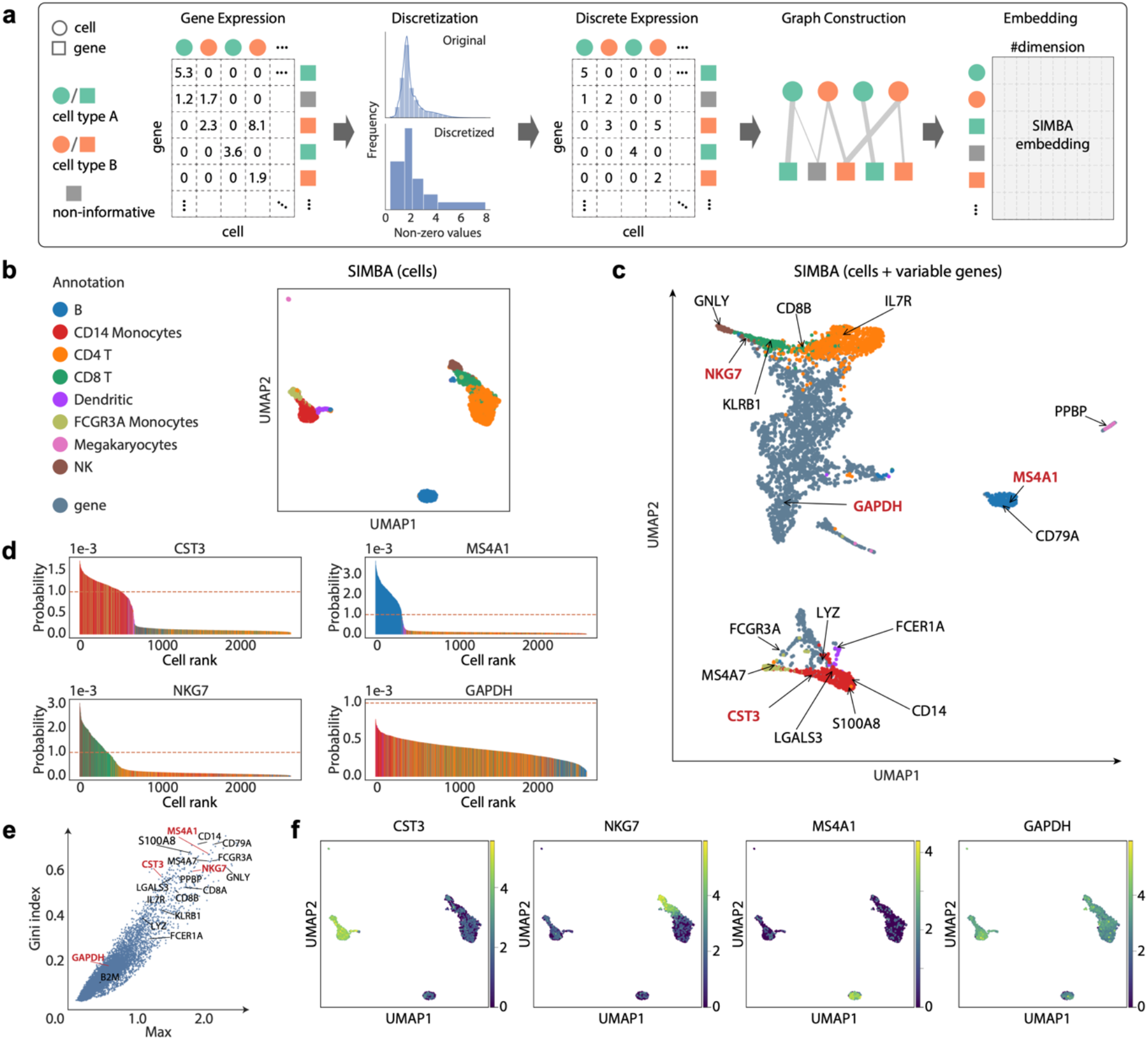
Single-cell RNA-seq analysis of the 10x PBMCs dataset using SIMBA. **(a)** SIMBA graph construction and embedding in scRNA-seq analysis. Biological entities including cells and genes are represented as shapes and colored by relevant cell types (green and orange). Non-informative genes are colored dark grey. Gene expression measurements for each cell are organized into a cell-by-gene matrix. These normalized non-negative observed values undergo discretization into five gene expression levels. Cells and genes are then assembled into a graph with nodes representing cells and genes, and edges between them representing different gene expression levels. This graph may then be embedded into a lower dimensional space resulting in a #entities x #dimension (by default, 50) SIMBA embedding matrix. **(b)** UMAP visualization of SIMBA embeddings of cells colored by cell type. **(c)** UMAP visualization of SIMBA embeddings of cells and variable genes. Cells are colored according to cell type as defined in b. Genes are colored slate blue. Cell-type-specific marker genes and housekeeping genes recovered by Scanpy are indicated with text and arrows. Genes highlighted in red are shown in **d**, **e**, and **f**. **(d)** SIMBA barcode plots of genes *CST3, MS4A1, NKG7*, and *GAPDH*. The x-axis indicates the ordering of a cell as ranked by the probability for each cell to be associated with a given gene. The y-axis describes the probability. The sum of probability over all cells is equal to 1. Each cell is one bar and colored according to cell type as defined in b. **€** SIMBA ranking of genes based on the proposed metrics. All the genes are plotted according to the Gini index against max score. The same set of genes as in **c** are annotated. **(f)** UMAP visualization of SIMBA embeddings of cells colored by gene expression of (left to right): *CST3, NKG7, MS4A1*, and *GAPDH*.

When the SIMBA embeddings of cells were visualized, each of the eight cell types, including B cells, megakaryocytes, CD14 monocytes, FCGR3A monocytes, dendritic cells, NK cells, CD4 T, and CD8 T cells, was clearly separated (**Fig. 2b**). When the SIMBA embeddings of both cells and genes were visualized, the co-embedding space showed that SIMBA not only recovered the cellular heterogeneity, but also correctly embedded informative genes close to relevant cell types (**Fig. 2c**). The same set of marker genes used to annotate these cells from Scanpy^2^ was highlighted on the UMAP plot. In addition, as a control, we also show the locations of two housekeeping genes *GAPDH* and *B2M*, which would not be expected to associate with any particular cell type. From the UMAP plot, we can see that SIMBA not only was able to embed major-cell-group specific genes to the correct locations (e.g., *IL7R* was embedded into CD4T cells and *MS4A1* was embedded into B cells), but also was robust to rare-cell-group specific genes (e.g., *PPBP* was embedded into megakaryocytes). On the contrary, non-informative or non-cell-type specific genes such as *GAPDH* and *B2M* were embedded in the middle of all cell groups (**Fig. 2c and Supplementary Fig. 1c**).

These highlighted genes can be further confirmed with “barcode plot”, which visualizes the estimated probability of assigning a feature to a cell by SIMBA based on the recovered edge confidence (**Fig. 2d, Supplementary Fig. 1e, Methods**). An imbalance in probability indicates the association of a gene to a sub-population of cells (often corresponding to known cell-types), whereas a uniform probability distribution indicates a non-cell-type-specific gene. For marker genes (*CST3* for monocytes and dendritic cells, *MS4A1* for B cells, and *NGK7* for NK and CD8T cells), we observed a clear excess in the probability of assigning each gene to their respective cell types.-Conversely, for the housekeeping gene *GAPDH*, we observed a more uniform distribution with much lower probability of associating that gene with the top-ranked cells.

SIMBA also provides several quantitative metrics (termed “SIMBA metrics”), including max value, Gini index, standard deviation, and entropy, to assess cell-type specificity of various features without requiring the prior knowledge such as cluster labels, predefined cell types, or known marker genes (**Methods**). As an example, by inspecting the gene metric plot of max value (a measurement of maximum probability, a higher value indicates higher cell-type specificity) vs Gini index (a measurement of imbalance, a higher value indicates higher cell-type specificity), we observed that the marker genes (e.g., *CST3, NKG7, MS4A1*) fall in the upper right corner, as opposed to housekeeping genes (e.g., *GAPDH*) in the lower left corner (**Fig. 2e**). Similar separation is observed with other metrics (**Supplementary Fig. 1b**). The cell type specificity of the selected marker genes was further confirmed by visualizing their expression pattern on UMAP plots (**Fig. 2f and Supplementary Fig. 1d**), accompanied by SIMBA barcode plots (**Supplementary Fig. 1d**). As a certain feature (e.g., genes) might notably outnumber cells or other features (when multiple types of features are present), SIMBA metrics not only serve as an efficient way of ranking features based on their cell type specificity, but also provides a straightforward way to filter out non-informative (non-cell-type-specific) features so that only the embeddings of cells and informative features will be visualized and the SIMBA space will not be crowded with non-informative features (e.g., house-keeping genes).

We next compared the top 600 marker genes identified by SIMBA (based on max value and Gini index) with those identified by the clustering-based statistical-tests method implemented in Scanpy (based on z-score calculated from the two-sided Wilcoxon rank-sum test with a Benjamini-Hochberg p-value correction, one of the statistical tests recommended in Scanpy’s tutorial) (**Supplementary Fig. 2a**). Upon comparison, we observed that nearly half of the marker genes discovered by SIMBA overlap with the marker genes identified by Scanpy (**Supplementary Fig. 2a**). However, on inspection of the top non-overlapping marker genes, genes identified by SIMBA are found to be enriched only within certain groups of cells (**Supplementary Figs. 2b and 2c**) while genes identified by Scanpy but not by SIMBA include the housekeeping gene *B2M* and multiple ribosomal protein genes (e.g., *RPS3* and *RPS6*) that are expressed ubiquitously in all cell types (**Supplementary Figs. 2b and 2d**). Furthermore, a combination of different statistical tests proposed in Scanpy is required to recover the genes identified only by SIMBA. For example, IL7R was identified only by using the t-test and FCER1A was identified only by using the Wilcoxon rank-sum test, as also noted in the Scanpy’s tutorial, while SIMBA successfully identified both *IL7R* and *FCER1A* as informative genes with a single procedure and without clustering the cells (**Fig. 2e and Supplementary Fig. 1b**). These examples illustrated some limitations of the clustering-based statistical-tests methods.

Lastly, we showed that SIMBA does not require variable gene selection, which is an essential step in standard scRNA-seq pipelines such as Seurat or Scanpy. SIMBA produces very similar embeddings for cells with and without variable gene selection (**Fig. 2b** and **Supplementary Fig. 2e**), though we observed that variable gene section does improve efficiency of the training procedure.

### SIMBA enables simultaneous characterization of cell states and cis-regulatory elements by jointly modeling accessible sites and DNA sequences in scATAC-seq analysis

As one of the most popular single-cell epigenomic techniques, single-cell assay for transposase-accessible chromatin using sequencing (scATAC-seq) has been widely used to profile regions of open chromatin and identify functional *cis*-regulatory elements such as enhancers and active promoters. In scATAC-seq, cells are characterized by different types of features ^14^, such as regions of accessible chromatin (“peaks” or “bins”) and *cis*-regulatory elements (DNA sequences) within these accessible regions including transcription factor (TF) motifs or *k*-mers.

Unlike existing methods that can only use peaks/bins or the DNA sequence within them, SIMBA can leverage simultaneously both types of features to learn cell states due to its flexibility in graph construction. Also, as SIMBA encodes cell-feature or feature-feature relations into the graph based on the simple binary presence of a feature, SIMBA does not need additional normalization steps such as term frequency-inverse document frequency (TF-IDF), which is required by most scATAC-seq analyses. When only peaks/bins are used, SIMBA constructs a graph with nodes representing cells and chromatin regions (peaks or bins) and edges indicating the accessibility of the chromatin regions in cells (**Fig. 3a**). When the DNA sequences for chromatin regions are available, SIMBA can also encode DNA sequences including TF motifs and *k*-mers into the graph by adding edges between these entities as nodes and the existing chromatin region nodes. The edges in this case indicate the presence of TF motifs/*k*-mers within these chromatin-accessible regions. Through the embedding procedure, SIMBA generates embeddings of cells along with peaks and DNA sequences (**Methods**). Finally, either the partial SIMBA embeddings (embeddings of cells in **Fig.3b**) or the whole SIMBA embeddings (embeddings of cells and all the features in **Fig.3c**) can be visualized. Therefore, SIMBA enables dimensionality reduction by leveraging both chromatin accessible regions and cis-regulatory sequences. Simultaneously, it highlights the cell-type-specific open chromatin regions and regulatory DNA sequences in a clustering-free way.

**Figure 3.**
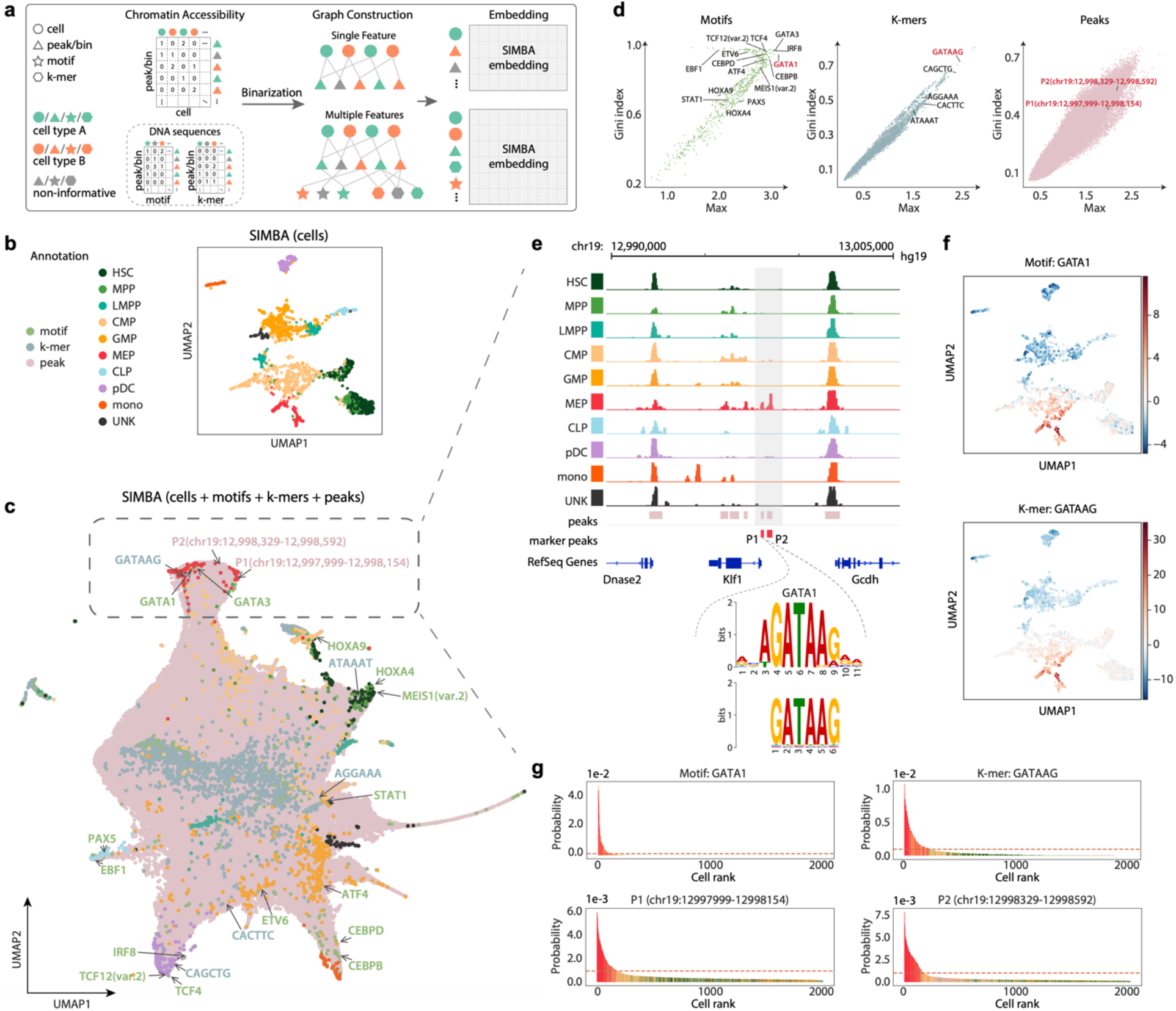
Single-cell ATAC-seq analysis of the human hematopoiesis dataset using SIMBA. **(a)** SIMBA graph construction and embedding in scATAC-seq analysis. Biological entities including cells, peaks/bins, TF motifs, k-mers are represented as shapes and colored by relevant cell types (green and orange). Non-informative features are colored dark grey. Cells and chromatin accessible features (peaks / bins) are organized into a cell x peaks / bins matrix. When sequence information (TF motif or *k*-mer sequence) within these regions is available, they can be organized into two sub-matrices to associate a TF motif or k-mer sequence with each peak/bin. These constructed feature matrices are then binarized and assembled into a graph. When a single feature (chromatin accessibility) is used, the graph encodes cells and peaks/bins as nodes. When multiple features (both chromatin accessibility and DNA sequences) are used, this graph may then be extended with the addition of TF motifs and k-mer sequences as nodes connected. Finally, SIMBA embeddings of these entities are generated through a graph embedding procedure. **(b)** UMAP visualization of SIMBA embeddings of cells colored by cell type. **(c)** UMAP visualization of SIMBA embeddings of cells and features including TF motifs, k-mers, and peaks. Cells are colored by cell type while motifs, k-mers, and peaks are colored green, blue, and pink, respectively. Cell type specific features that are embedded near their corresponding cell types are indicated as the text labels (colored according to feature type) with arrows. **(d)** SIMBA metric plots of TF motifs, k-mers, and peaks. Cell-type specific features annotated in **(c)** are highlighted. **€** Genomic tracks of aligned scATAC-seq fragments, separated and colored by cell type. Two marker peaks P1 and P2 in red are shown beneath the alignment. Within the peak P1, k-mer GATAAG and its resembling GATA1 motif logo are highlighted. **(f)** UMAP visualization of SIMBA embeddings of cells colored by TF activity scores of the GATA1 motif and k-mer GATAAG enrichment. **(g)** SIMBA barcode plots of the GATA1 motif, k-mer GATAAG, and the two peaks P1 and P2. Cells are colored according to cell type labels described above. Dotted red line indicates the same cutoff used in all four plots.

To demonstrate the value of SIMBA embeddings for scATAC-seq analysis, we first applied SIMBA to a scATAC-seq data of 2,034 human hematopoietic cells with FACS-characterized cell types^15^(**Supplementary Table 1**). For the SIMBA embeddings of cells alone, as shown in **Fig. 3b**, SIMBA accurately separated cells such that cells belonging to distinct cell types are visually separated. For the SIMBA embeddings of cells together with various types of features, as shown in **Fig. 3c**, SIMBA successfully embedded distinct features from both positional (peaks/bins) as well as sequence-content (TF motifs and k-mers) information together based on their biological relations. Notably, based on SIMBA metrics, these highlighted features that are embedded within each cell type all have high cell-type specificity scores (shown in the upper right part of SIMBA metric plots in **Figure 3d**).

Our analysis using SIMBA led to several key findings in human hematopoietic differentiation.

First, SIMBA identified key master regulators of hematopoiesis. As highlighted in **Fig. 3c**, we observed that motifs of previously reported TFs were embedded near their respective cell types in the UMAP plot. For example, the GATA1 and GATA3 motifs are proximal to megakaryocyte-erythroid progenitor (MEP) cells^16^, the PAX5 and EBF1 motifs are near to common lymphoid progenitor (CLP) cells^17^, and the CEBPB and CEBPD motifs are proximal to monocyte (mono) population^18^.

Second, SIMBA revealed an unbiased set of DNA sequences, i.e., *k*-mers, that represent important TF binding motifs involved in hematopoiesis. We observed that these *k*-mers were embedded near their resembling TF binding motifs and relevant cell subpopulations (**Fig. 3c** and **3e**, **Supplementary Fig. 3b**), indicating that this methodological framework is capable of *de novo* motif discovery. For example, the DNA sequence, CAGCTG is embedded in plasmacytoid dendritic cells (pDCs); this sequence matches the TCF12 binding motif, which controls dendritic cell lineage specification. To further illustrate the interpretability of the SIMBA embeddings of TF motifs and k-mers, we calculated per-cell TF activity scores^19^ (high-variance TF motifs/*k*-mers) and visualized them on SIMBA embeddings of cells. As shown in **Figure 3f**, the GATA1 TF motif and k-mer GATAAG that were both embedded in MEP cells by SIMBA, also showed high-level activity in MEP cells. The consistency between SIMBA embedding and TF activity was observed for most of other TF motifs and *k*-mers as well (**Supplementary Fig. 3a, 3b**).

Third, SIMBA identified differentially accessible chromatin regions that may mediate cell-type specific gene regulation. For example, the two peaks with coordinates chr19:12997999-12998154 (P1) and chr19:12998329-12998592 (P2) that were embedded within MEP cells were almost exclusively observed in MEP cells on KLF1 genome track (**Fig. 3e**). Interestingly, P1, upstream of *KLF1*, contains the *k*-mer GATAAG that matches the GATA1 binding motif, while transcription factor GATA1 is known to regulate the gene *KLF1* and plays a pivotal role in erythroid cell and megakaryocyte development^20^. Therefore, by embedding these MEP-cell-related regulatory elements into the neighborhood of MEP cells, SIMBA demonstrates a novel means of studying the epigenetic landscape of cell differentiation. To further validate the differentially accessible regions identified by SIMBA, we selected 100 peaks at random from each annotated cell type in SIMBA co-embedding space. From the heatmap of chromatin accessibility, we clearly see that the peaks embedded nearby respective types correlate with strong cell-type specificity. This observation is robust to the number of cells within each cell type (**Supplementary Fig. 3c**).

Available methods for scATAC-seq analysis visualize only cells. While SIMBA diverges from these available workflows, enabling the co-embedding of cells and features, we still qualitatively and quantitatively compared the SIMBA embeddings of cells to state-of-the-art scATAC-seq analysis methods by their ability to distinguish cell types. Our analyses show that SIMBA overall performs better than the methods evaluated, further demonstrating the wide utility of SIMBA (**Supplementary Figs. 4 and 5; Supplementary Note 1**).

### SIMBA enables simultaneous learning of cellular heterogeneity and gene regulatory circuits from integrated analysis of single-cell multimodal data

scRNA-seq and scATAC-seq are two of the most widely adopted single-cell sequencing technologies, but they are limited to measuring only a single aspect of cell state at a time. To improve our ability to interrogate cell states, several single-cell dual-omics technologies have been recently developed ^21–24^ to jointly profile transcriptome and chromatin accessibility within the same individual cells, therefore providing the potential to correlate gene expression with accessible regulatory elements and further delineate the yet elusive principles of gene regulation. This section outlines the SIMBA’s ability to simultaneously learn cell heterogeneity as well as gene regulatory circuits from single-cell multiomic data. We applied SIMBA to three recent single-cell dual-omics technologies: SHARE-seq^22^, SNARE-seq^21^, and a multiome PBMCs dataset from 10x Genomics (**Supplementary Table 1**).

**Figure 4a** illustrates the procedure of graph construction and generation of the final SIMBA embedding matrix. Briefly, for scRNA-seq, the gene expression matrix is discretized to generate different levels of gene expression. For scATAC-seq, both the chromatin accessibility matrix and motif/*k*-mer match matrix are binarized. In this graph, there are five entity (node) types, including cells, genes, peaks, motifs, and *k*-mers. For scRNA-seq, an edge indicates whether a gene is expressed in a cell and its weight indicates the gene expression level (five levels, by default). For scATAC-seq, an edge indicates whether a peak is present in a cell or if a TF motif/*k*-mer is present within a peak. Once the graph is constructed, the graph embedding procedure is performed to generate SIMBA embeddings of cells and different types of features. scATAC-seq peaks generally greatly outnumber cells and other features and many of these peaks are non-informative, resulting in them dominating the space if the whole SIMBA embeddings are visualized (**Supplementary Fig. 6a, c**). In such cases, we leverage the flexibility of SIMBA embedding to only visualize the partial SIMBA embeddings to improve the visibility of cells and cell-type-specific features.

**Figure 4.**
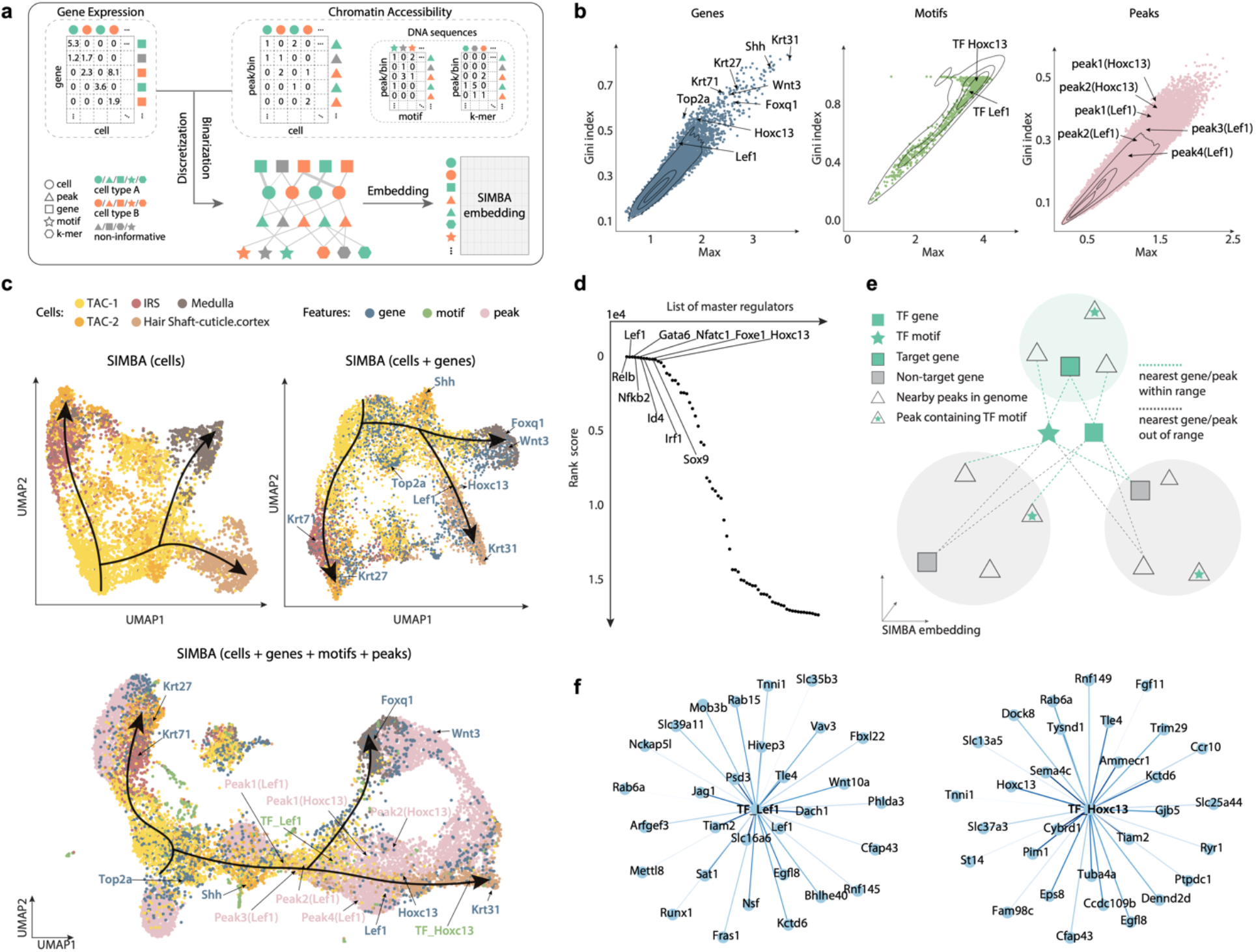
Multimodal analysis of the SHARE-seq hair follicle dataset using SIMBA. **(a)** SIMBA graph construction and embedding in multimodal analysis. Overview of SIMBA’s approach to multimodal (scRNA-seq + scATAC-seq) data analysis. **(b)** SIMBA metric plots of genes, TF motifs, and peaks. All these features are plotted according to the Gini index against max score. Cell-type specific genes, TF motifs, and peaks are highlighted. **(c)** UMAP visualization of SIMBA embeddings of cells (Top-left), cells and genes (Top-right), and cells along with genes, TF motifs, and peaks (Bottom). **(d)** Ranked scatter plot of candidate master regulators as identified by SIMBA. **€** Schematic description of SIMBA’s strategy for identifying target genes given a master regulator. **(f)** Top 30 target genes of transcription factors Lef1 and Hoxc13 as inferred by SIMBA. To demonstrate the usefulness and versatility of the SIMBA embeddings, we analyzed the cell populations undergoing the dynamic process of hair follicle differentiation from mouse skin profiled with SHARE-seq.

First, we calculated SIMBA metrics (max values and Gini index scores) to assess the cell-type specificity of different types of features, including genes, TF motifs, and peaks (**Fig. 4b, Methods**). As shown in **Figure 4b**, based on these metrics, we successfully recovered genes associated with hair follicles such as *Lef1* and *Hoxc13*. Similarly, TF motifs and peaks proximal to the genomic loci of these genes also score in the upper right quadrant of the metric plots. SIMBA’s cell-type specificity metrics successfully revealed the key genes and regulatory factors important to the hair follicle differentiation process.

Next, we visualized and interrogated the SIMBA embeddings of 1) cells; 2) cells and top-ranked genes based on SIMBA metrics; and 3) cells, top-ranked genes and TF motifs based on SIMBA metrics, and the neighboring peaks of these genes and TF motifs by querying the SIMBA space (**Methods**). **Figure 4c** shows the UMAP visualization of the partial SIMBA embeddings of cells and informative features. The UMAP visualization of SIMBA embeddings of cells and the full set of features was also performed (**Supplementary Fig. 6a**).

The SIMBA embeddings of cells were able to reveal the three fate decisions from transit-amplifying cells (TACs), including inner root sheath (IRS), medulla, and cuticle/cortex. The SIMBA embeddings of cells and informative features uncovered important genes and regulatory factors along the hair follicle differentiation trajectories. For example, the marker genes *Krt71, Krt31*, and *Foxq1* were embedded into their corresponding cell types: IRS, cuticle/cortex, and medulla, respectively. The Lef1 motif was embedded into the beginning of medulla and cuticle/cortex lineages while the Hoxc13 motif was embedded into the late stage of cuticle/cortex differentiation. Peaks near the *Lef1* and *Hoxc13* loci were also embedded into the nearby regions of these genes and motifs, as expected.

To show the robustness of SIMBA, we separated the scRNA-seq and scATAC-seq components within the SHARE-seq dataset and performed each respective single-modality analysis. With the consistent embedding results of cells as in multimodal analysis, we further demonstrated that SIMBA embedding procedure is robust to the type and the number of features encoded in the input graph (**Supplementary Fig. 6b**,**6c**). Each reported marker gene was corroborated using the UMAP plots with cells colored by gene expression as well as using the SIMBA barcode plots. The two aforementioned TF motifs and their respective peak sets were supported by the corresponding SIMBA barcode plots, wherein we observed an imbalanced distribution with high probability towards the correct cell type labels (**Supplementary Fig. 7a-d**).

Further, we demonstrated that the SIMBA co-embedding space of cells and features provides the potential to identify master regulators of differentiation and infer their target regulatory genes. To define a master regulator *a priori*, we postulated that both its TF motif and TF gene should be cell-type specific, given that active gene regulation involves both the expression of a TF and accessibility of its binding sites. Thus, the TF motif and TF gene should be embedded closely in the shared latent space. Extending this logic to identify putative master regulators, we assessed the cell-type-specificity of TF motifs and genes based on SIMBA metrics and ranked all potential master regulators based on the distance between the TF motif and the respective TF gene in the shared SIMBA embedding space (**Methods**). SIMBA successfully identified previously described master regulators such as Lef1, Gata6, Nfatc1, and Hoxc13 as the top master regulators related to lineage commitment in mouse skin (**Fig. 4d, Supplementary Table 2**). To infer the target genes of a given master regulator, we postulate that in the shared SIMBA embedding space, 1) the target gene is close to both the TF motif and the TF gene; 2) the accessible regions (peaks) near the target gene loci must be close to both the TF motif and the target TF gene. Resting on these assumptions of *cis*-regulatory dynamics, the inference of target genes was performed by calculating the distance between target gene candidates and the respective TF motif and gene. In addition, nearby peaks around the target gene’s locus and the presence of TF motif in these nearby peaks are also considered (**Fig. 4e, Methods**). The top 30 target genes of TF Lef1 and TF Hoxc13 inferred by SIMBA are shown respectively (**Fig. 4f, Supplementary Fig. 7e**). The full list of ranked target genes is provided in **Supplementary Table 3**. Notably, our approach recovered targets genes that were also reported in the original study^22^. For example, genes *Lef1, Jag1, Hoxc13, Gtf2ird1* are regulated by the TF Lef1, while genes *Cybrd1, Hoxc13, St14* are regulated by the TF Hoxc13.

In addition to SHARE-seq, we also applied SIMBA to another two dual-omics datasets, the mouse cerebral cortex dataset profiled by SNARE-seq^21^ (**Supplementary Fig. 8**) and the multiome PBMCs dataset from 10x Genomics (**Supplementary Fig. 9**). By validating the embeddings of cells and features with given cell type labels (**Supplementary Fig. 8a and Fig.9a**), marker genes from the original study (**Supplementary Fig. 8a,b,d and Fig. 9a,b,d**), and differentially accessible chromatin regions (**Supplementary Fig. 8c and Fig. 9c**), we further demonstrate the suitability of SIMBA for multimodal analysis.

### SIMBA enables simultaneous batch correction and clustering-free marker gene detection

Efforts to collect data from single cells has grown to the level of consortia that span multiple institutions with the hopes of finely mapping and characterizing specific tissues. This has brought with it an increased demand for analysis methods that are capable of negating technical covariates inherent to multi-batch data collection, including experimental replicate identity, sample preparation, and sequencing platform. Batch correction that removes the effects of technical covariation while preserving true biological signals is required prior to downstream analysis ^25, 26^. Existing methods follow a workflow with four primary steps. The first step is the actual batch correction, which often generates a “batch corrected” latent space. The second step clusters cells in this batch corrected space. Based on the clustering result the third step detects marker genes in the original gene expression space of each batch because the low-dimensional “batch corrected” space is no longer comprised of genes. The fourth step finally combines the marker genes detected from each batch. However, these methods are clustering-dependent and may result in the inconsistent explanation of marker genes as marker genes are detected in each original batch as opposed to the batch-corrected space. Unlike current methods, in addition to embeddings of cells, SIMBA generates comparable embeddings of genes and therefore relieves marker gene discovery from a dependence on the original gene expression space. Thus, SIMBA enables simultaneous batch effect removal and cell-type-specific marker gene detection in the same integrated space without clustering.

We first demonstrate that SIMBA readily corrects batch effects and produces joint embeddings of cells and genes across multiple scRNA-seq datasets generated from varying sequencing platforms and cell type compositions. While existing methods for scRNA-seq analysis rely on specialized tools for batch correction, SIMBA works as a stand-alone method obviating the need for prior input data correction when applied to multi-batch scRNA-seq dataset. SIMBA accomplishes batch correction by encoding multiple scRNA-seq datasets into a single graph (**Fig. 5a**). Cell nodes across batches are connected to gene nodes through experimentally measured edges as in the previously described scRNA-seq graph construction. Here, the gene nodes are shared between the cell nodes of different batches. In addition to the experimentally measured edges, batch correction is further enhanced through computationally inferred edges drawn between similar cell nodes across datasets using a truncated randomized singular value decomposition (SVD)-based procedure. SIMBA then generates the embeddings of all nodes including cells of each batch and genes from the resulting graph (**Methods**). The SIMBA embeddings of cells naturally represent the batch-corrected space. In addition, the whole SIMBA embeddings of all entities provide the batch-corrected space, in which cells and genes co-exist, and therefore allow for individual-cell-level marker detection by performing biological queries of cells in the SIMBA space (**Methods**). We visualized both SIMBA embeddings of cells (**Fig. 5b**), and the whole SIMBA embeddings of cells and genes (**Fig. 5c**) in UMAP.

**Figure 5.**
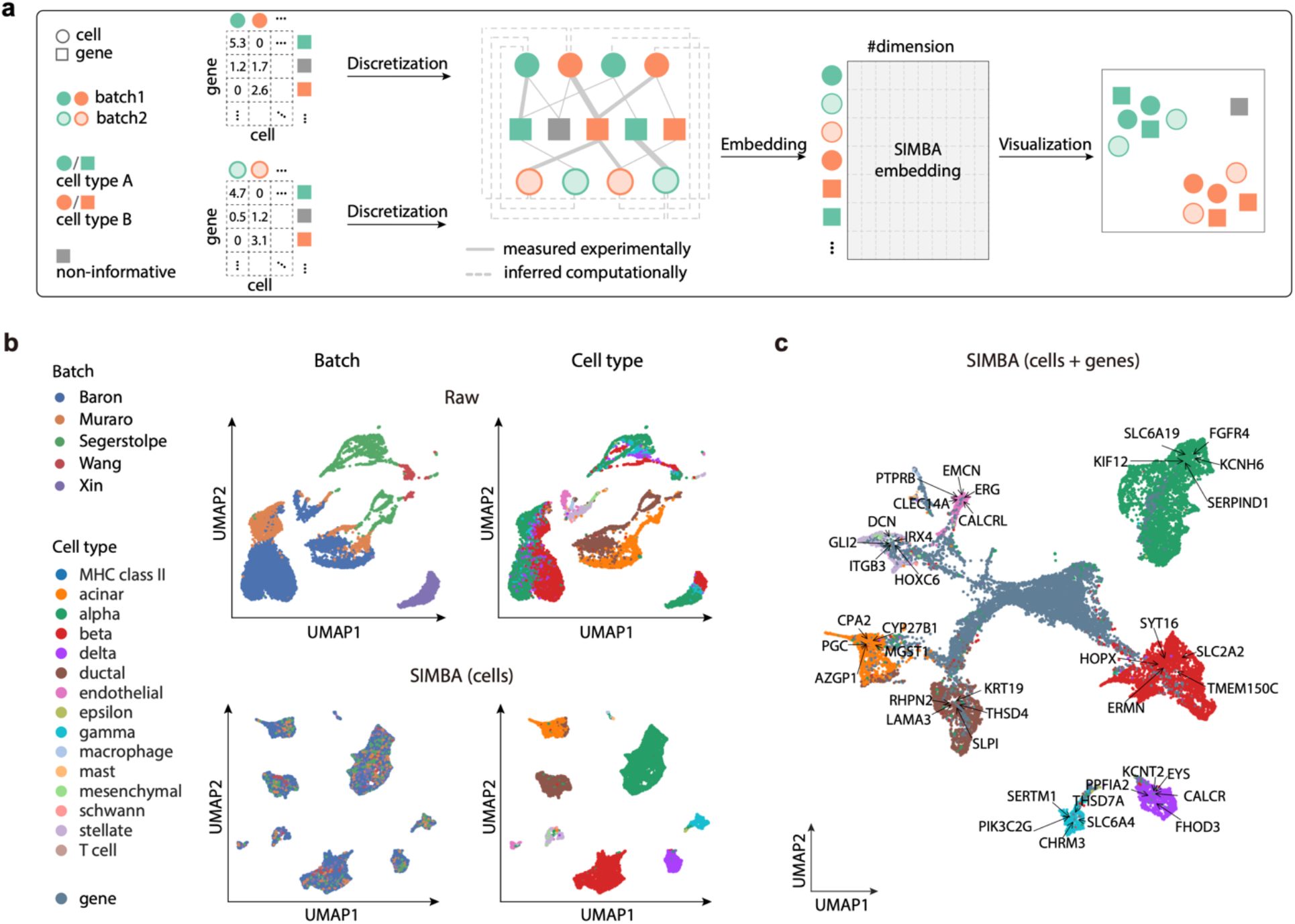
Batch correction analysis of scRNA-seq data using SIMBA. **(a)** SIMBA graph construction and embedding in batch correction analysis. Overview of SIMBA’s approach to batch correction across scRNA-seq datasets. Distinct shapes indicate the type of entity (cell or gene). Colors distinguish batches or cell types. **(b)** UMAP visualization of the scRNA-seq human pancreas dataset with five batches of different studies before and after batch correction. Cells are colored by scRNA-seq data source and cell type respectively. Top: UMAP visualization before batch correction; Bottom: UMAP visualization after batch correction with SIMBA; **(c)** UMAP visualization of SIMBA embeddings of cells and genes, with batch effect removed and known marker genes highlighted.

We applied SIMBA to two multi-batch scRNA-seq datasets; a mouse atlas dataset composed of two batches and a human pancreas dataset spanning five batches used in a recent benchmark study^25^ (**Supplementary Table 1**). The mouse atlas dataset contains two scRNA-seq datasets with shared cell types from different sequencing platform. The human pancreas dataset contains five samples pooled from five sources using four different sequencing techniques, in which not all cell types are shared across each sample. For both datasets, SIMBA successfully corrected batch effects, evenly mixing batches within annotated cell type clusters, while maintaining the segregation of these clusters in the resulting embedding, indicating preservation of biological signal and elimination of confounding technical covariates (**Fig. 5b, Supplementary Fig. 12b**). It is important to note that the mouse atlas dataset was collected from nine different organ systems, so there exists some expected heterogeneity within cell type labels. Conversely, the human pancreas datasets are curated from a single organ and SIMBA sufficiently separated cell types into transcriptionally distinct, homogeneous cell clusters (**Fig. 5b**).

Through removing batch effects during graph embedding, SIMBA simultaneously identifies cell-type-specific marker genes (**Fig. 5c**). In the absence of the eliminated technical covariation, marker genes are identifiable by performing biological queries for neighboring genes within cell types in the SIMBA embeddings of cells and genes (**Methods**). In the case of unknown cell labels, marker genes can be identified by calculating SIMBA metrics (**Methods**). SIMBA correctly embeds known cell-type-specific marker genes proximal to the correct cell type labels, while non-marker genes were non-proximal to specifically-labelled cells (**Supplementary Fig. 10, 11**). The resulting marker genes recapitulated the clustering-based differential expression (DE) analysis results for each dataset^27–32^ (e.g. *Cdh5, Tie1, Myct1* for endothelial cell and *C1qc*, Fcgr1 for macrophage, *S100a8, Trem3* for Neutrophil in the mouse atlas dataset and *KIF12* for alpha cell and *KRT19* for ductal cell in the human pancreas dataset) and are shown to be expressed specifically in the queried cell types (**Supplementary Fig. 10, 11**). Taken together, these results distinguish SIMBA from existing batch correction methods that rely on clustering in a batch-corrected space, followed by differential gene expression analysis in the original, uncorrected space of each batch.

While SIMBA is a versatile graph embedding method that can perform multiple tasks and generate embeddings of both cells and genes, we evaluated the SIMBA embeddings of cells for this task with methods that were specifically designed for batch correction. We considered three widely adopted batch correction methods that demonstrated top-tier performance based on a recent benchmark study^25^: Seurat3, LIGER and Harmony. Our results indicate that SIMBA achieved comparable batch correction performance both qualitatively and quantitatively while enabling simultaneous marker gene detection by providing the additional SIMBA embeddings of genes. (**Supplementary Note 2, Supplementary Figure 12**).

### SIMBA enables simultaneous multi-omics integration and clustering-free multi-type marker feature detection

Single-cell assays are now capable of measuring a broad range of cellular modalities and data is being generated that describes cells by varying features sets, which has motivated the need for methods that leverage these features to perform multi-omics integration such that a more comprehensive description of cell state may be learned. This is different from multi-modal analysis because the correspondence between individual cells is unknown. Current multi-omics integration methods follow a similar workflow as the previously described batch correction methods, including: 1) generating a low-dimensional integrated space of cells; 2) clustering cells in the integrated space; 3) detecting marker features in the original feature (e.g., genes, peaks) space of each modality because the low-dimensional integrated space no longer consists of the original features. Unlike existing multi-omics integration methods that cannot directly explore multi-type features in the integrated space and require clustering for identifying marker features, we demonstrate that SIMBA enables simultaneous multi-omics integration and clustering-free detection of distinct marker features, specifically as it is applied to datasets comprised of scRNA-seq and scATAC-seq.

SIMBA accomplishes this integration by first building one graph for scRNA-seq data and another graph for scATAC-seq data independently as described in previous sections (**Fig. 6a**). To connect these two graphs, SIMBA then calculates gene activity scores by summarizing accessible regions from scATAC-seq data and then infers edges between cells of different assays based on their shared gene expression modules as previously described in the batch correction section. Finally, SIMBA embeds the graph of cells, genes, and peaks into a common, low-dimensional space. The SIMBA embeddings of cells naturally represent the integrated space of multiple modalities. Furthermore, the SIMBA embeddings of all entities provide the integrated space containing cell, genes, and peaks, and therefore enable the individual-cell-level marker detection of multi-type features by performing biological queries of cells in SIMBA space (**Methods**). The SIMBA embeddings of these multi-omics entities can be visualized either partially or as a whole using UMAP or similar visualization tools.

**Figure 6.**
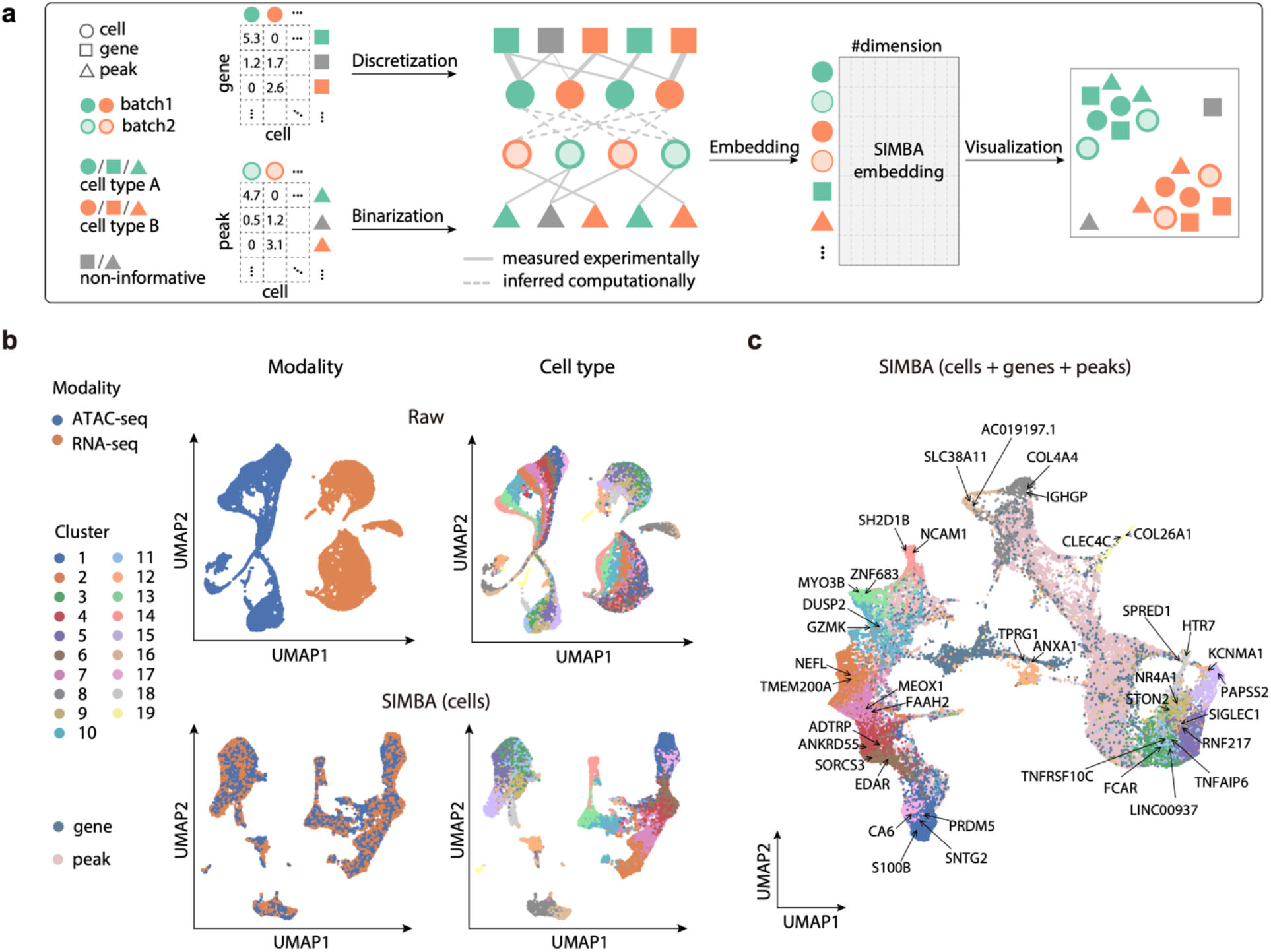
Multi-omics integration of scRNA-seq + scATAC-seq data using SIMBA. **(a)** SIMBA graph construction and embedding in multi-omics integration. Overview of SIMBA’s approach to data integration across scRNA-seq and scATAC-seq. Distinct shapes indicate the type of entity (cell, gene, or peak). Colors distinguish batches or cell types. **(b)** UMAP visualization of the integrated scRNA-seq and scATAC-seq data manually created from the 10x human PBMCs dataset before and after data integration. Cells are colored by single-cell modality and cell type respectively. Top: UMAP visualization before integration; Bottom: UMAP visualization after integration with SIMBA. **(c)** UMAP visualization of SIMBA embeddings of cells, genes, and peaks with two cell modalities integrated and known marker genes highlighted.

To facilitate the evaluation of data integration performance, we created datasets with ground-truth labels by manually splitting the dual-omics datasets into two single-modality datasets (i.e., scRNA-seq and scATAC-seq), in which we know the true matching between cells across the two modalities. We then applied SIMBA to the integration analysis of two case studies where scRNA-seq and scATAC-seq datasets are generated from the SHARE-seq mouse skin dataset and the 10x Genomics multiome human PBMCs dataset, respectively (**Supplementary Table 1**).

We first visualized the SIMBA embeddings of cells and observed that SIMBA was able to preserve cellular heterogeneity while evenly mixing the two modalities (**Fig. 6b, Supplementary Fig. 15b**). We then visualized the SIMBA embeddings of cells, genes, and top-ranked peaks based on SIMBA metrics and observed that in addition to learning cellular heterogeneity, SIMBA simultaneously identified marker genes and peaks at single-cell resolution. In the co-embedding space, we observed that the neighbor genes of cells (highlighted in UMAP plots), are each exclusively expressed in their corresponding cell types (**Supplementary Figs. 13a-e, 14a-c,e**). For example, in the SHARE-seq mouse skin dataset, *Foxq1* and *Shh* are located within medulla and TAC-2, respectively; in the 10x PBMCs dataset, *PAPSS2* and *KCNMA1*, which are the marker genes of blood monocytes, are embedded close to each other. Similarly, we observed that the neighbor peaks of cells show a clear cell-type-specific accessibility pattern that is robust to the cluster size of a given cell type (**Supplementary Figs. 13f and 14d).**

The joint embedding of cells and features produced by SIMBA is fundamentally distinguished from other multi-omics integration methods in that it simultaneously achieves integration as well as marker feature discovery. However, we still sought to compare the SIMBA embeddings of cells with two widely-adopted single-cell multi-omics integration methods, Seurat3 and LIGER, based on their ability to integrate single-cell modalities while persevering cellular heterogeneity (**Supplementary Note 3**). We observed that SIMBA achieved the overall best performance on the mouse skin SHARE-seq dataset and 10x PBMCs multiome dataset.

## Discussion

Multimodal measurements of individual cells offer new and unexplored opportunities to study cell identity as a function of the complex interactions between omic layers. While these datasets offer an exciting potential for discovery, computational analysis methods to fully delineate the cell states and molecular processes across multiple genomic features remain insufficient.

As presented in this manuscript, SIMBA models cells and measured features as nodes encoded in a graph and employs a scalable and efficient graph embedding procedure to embed nodes of cells and features into a shared latent space. We demonstrate that direct graph representations of single-cell data capture not only the relations between cells and the quantified features of the experiment (e.g., gene expression or chromatin accessibility) but also hierarchical relations between features. An example of such a hierarchical relation is the coordinate-level description of an ATAC-seq peak and the corresponding TF motifs and/or *k*-mer sequences contained within that region. In the resulting joint embedding, proximity-based biological queries can be performed to discover cell-type-specific co-regulatory machinery across modalities. Therefore, SIMBA enables simultaneous learning of cellular heterogeneity and cell-type-specific multimodal features and complements the current gene regulatory network analyses. SIMBA also circumvents the ordinary reliance on cell clustering for cell sub-population feature discovery. We thus avoid user-defined clustering solutions, which may lead to artifactual discovery or false negative results.

SIMBA has been extensively benchmarked across single-cell modalities and tasks, obtaining better or comparable performance metrics when compared to current state-of-the-art methods developed for the respective task. In contrast to tools developed and optimized for a single, specific task these results suggest a wide applicability of SIMBA’s graph-based framework, obviating the need to combine multiple analysis tools.

Graph embedding methods hold significant promise for the analysis of biological data. Previous applications of graph embedding include functional annotation of genes ^33^, transcription factor binding to DNA motifs ^34^ and more recent single-cell RNA-seq analyses ^35, 36^. The graph encoding and embedding procedures we have outlined may be potentially improved and extended to better represent biological entities and capture their respective relations.

Foreseeable extensions of SIMBA may include the analyses of increasingly complex datasets. For example, in the analysis of spatial transcriptomics wherein transcriptomic measurements are mapped to the true cell coordinates within a tissue ^37^, we can encode the spatial proximity into a SIMBA graph. We also envision extending this framework to data describing 3-D chromatin conformation wherein the interaction between DNA segments can be encoded to represent how regulatory regions are linked to genes^38^. Another potential extension of SIMBA could consider single-cell lineage-tracing datasets^39^ wherein both cellular lineage information and gene expression measurements are captured and can be potentially encoded into a SIMBA graph to represent their longitudinal relations. In general, we are interested in the further incorporation of external information and hierarchical relations between features in the graph. We anticipate our comprehensive and extensible SIMBA framework (https://simba-bio.readthedocs.io/) will provide the possibility to leverage *a priori* knowledge for graph embedding and the flexibility to extend to new experimental designs.

It is likely that multi-omics assays will continue to improve as well as expand in scope. Already, innovation in these data-generating technologies have outpaced the development of corresponding computational frameworks required to gain integrative insights from such rich data. This disparity highlights a need for methods that break through previous limitations and are easily extended to future cell measurements. We believe SIMBA satisfies these conditions as a comprehensive and extensible method for exploring cellular heterogeneity and investigating the regulatory mechanisms that drive cellular diversity while laying a groundwork for the development of new non-cluster-centric analysis methods for single cell omics data.

## Methods

### Single-cell data preprocessing

#### a. Single-cell RNA-seq

Genes expressed in fewer than three cells were filtered. Raw counts were library size-normalized and subsequently log-transformed. Optionally, variable gene selection ^12^ (a python version is implemented in SIMBA that is inspired by Scanpy^2^) may be performed to remove non-informative genes and accelerate the training procedure. Notable differences in the resulting cell embeddings were not observed upon limiting feature input to those identified by variable gene selection but SIMBA embeddings of non-variable genes will not be generated as they are not encoded in the graph.

#### b. Single-cell ATAC-seq

Peaks present in fewer than three cells were filtered. Optionally, we implemented a scalable truncated-SVD-based procedure to select variable peaks as a preliminary step to additionally filter non-informative peaks and accelerate the training procedure. First the top k principal components (PCs) were selected, with k chosen based on the elbow plot of variance ratio. Then for each of the top *k* PCs, peaks were automatically selected based on the loadings using a knee point detection algorithm implemented by ‘kneed’^40^. Finally, peaks selected for each PC were combined and denoted as “variable peaks”. Similar to the observation made with scRNA-seq data, the optional step of variable peak selection has a negligible effect on the resulting cell embedding. Despite this minimal impact on the resulting embedding, this feature selection step imparts a significant practical advantage in reducing training procedure time.

*k*-mer and motif scanning was performed using packages ‘Biostrings’ and ‘motifmatchr’ with JASPAR2020^41^. Included in the implementation of SIMBA is a convenient R command line script “scan_for_kmers_motifs.R”, which will convert a list of peaks (formatted in a bed file) to a sparse peaks-by-*k*-mers/motifs matrix, which is stored as an hdf5-formated file.

### Graph construction (five scenarios)

#### i. Single-cell RNA-seq analysis

The distribution of non-zero values in the normalized gene expression matrix was first approximated using a *k*-means clustering-based procedure. First, the continuous non-zero values were binned into *n* intervals (by default n=5). Bin widths were defined using 1-dimensional *k*-means clustering wherein the values in each bin are assigned to the same cluster center. The continuous matrix is then converted into a discrete matrix wherein1, …, *n* are used to denote *n* levels of gene expression. Zero values are retained in this matrix. Then the graph was constructed by encoding two types of entities, cells and genes, as nodes and relations with *n* different weights between them, i.e., *n* levels of gene expression, as edges. These *n* relation weights range from 1.0 to 5.0 with a step size of 5/n denoting gene expression levels (lowest: 1.0, highest: 5.0), such that edges corresponding to high expression levels affect embeddings more strongly than those with intermediate or low expression levels. This discretization is implemented in the SIMBA package using the function, “si.tl.discretize()”.

#### ii. Single-cell ATAC-seq analysis

Peak-by-cell matrices were binarized, with “1” indicating at least one read within a peak and “0” otherwise. The graph was constructed by encoding two types of entities, cells and peaks, as nodes and the relation between them, denoting the presence of a given peak in a cell, as edges. The single relation type was assigned with a weight of 1.0. When the DNA sequence features were available, they were encoded into the graph using *k*-mer and motif sequence entities as nodes. This was performed by first binarizing the peak-by-*k*-mer/motif matrix then constructing an extension to the original peak/cell graph using the peaks, *k*-mers, and motifs as nodes and the presence of these entities within peaks as edges between these additional nodes and the peak nodes. The relation between k-mers and peaks was assigned a weight of 0.02 while the relation between TF motifs was assigned a weight of 0.2. Of note, *k*-mers and motifs may be used independently of each other as node inputs to the graph, depending-on the specific analysis task.

#### iii. Multimodal analysis

Combination of the above outlined strategies for graph construction of scRNA-seq and scATAC-seq data was used to construct a multi-omics graph.

#### iv. Batch correction

A graph for each batch was constructed as described in i). Edges between cells of different batches were inferred through a procedure based on truncated randomized singular value decomposition (SVD) to link disjoint graphs of different batches. More specifically, in the case of scRNA-seq data, consider two gene expression matrices *X*1_*n*_1_×*m*_ and *X*2_*n*_2_×*m*_, where *n*_1_, *n*_2_ denotes the number of cells and *m* denotes the number of the shared features, i.e., variable genes, between datasets. The matrix *X*_*n*_1_×*n*_2__ was then computed by multiplying *X*1 and *X*2:

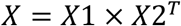

Truncated randomized SVD was subsequently performed on *X*:

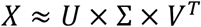

where *U* is an *n*_1_ × *d* matrix, Σ is an *d* × *d* matrix, and *V* is an *n*_2_ × *d* matrix (by default *d* = 20).

Both *U* and *V* were further *L*2 normalized. For each cell in *U*, we searched for *k* nearest neighbors in *V* and vice versa (by default, *k* = 20). Eventually, only the mutual nearest neighbors between *U* and *V* were retained as inferred edges between cells (represented as dashed lines in **Fig. 5a**). The procedure of inferring edges between cells of different batches is implemented in the function “si.tl.infer_edges()” in the SIMBA package.

For multiple batches, SIMBA can flexibly infer edges between any pair of datasets. In practice, however edges are inferred between the largest dataset(s) or the dataset(s) containing the most complete set of expected cell types and other datasets.

#### v. Multi-omics integration

scRNA-seq and scATAC-seq graphs were constructed following steps i) and ii), respectively. To infer the edges between cells of scRNA-seq and scATAC-seq, gene activity scores were first calculated for scATAC-seq data^3^. More specifically, for each gene, peaks within 100kb upstream and downstream of the TSS were considered. Peaks overlapping gene body region or within 5kb upstream of gene bodies were given the weight of 1.0. Otherwise, peaks were weighted based on their distances to TSS using the exponential decay function: 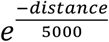. Subsequently, the gene score of each gene was computed as a weighted sum of the considered peaks. These gene scores were then scaled to respective gene size. These steps are implemented by the function “si.tl.gene_scores()” in SIMBA. For user convenience, the SIMBA package curates the gene annotations of several commonly used reference genomes, including hg19, hg38, mm9, and mm10. Once gene scores were obtained, the same procedure described in iv) was performed to infer edges between cells profiled by scRNA-seq and scATAC-seq using the function, “si.tl.infer_edges()” in SIMBA.

The procedure of generating constructed graphs is implemented in the function, “si.tl.gen_graph()” in the SIMBA package.

### Graph Embeddings with Type Constraints

Following the construction of a multi-relational graph between biological entities, we adapted graph embedding techniques from the knowledge graph and recommendation systems literature to construct unsupervised representations for these entities.

We provide as input a directed graph *G* = (*V, E*), where *V* is a set of entities (vertices) and *E* is a set of edges, with a generic edge *e* = (*u, v*) between a source entity *u* and destination entity *v*. We further assume that each entity has a distinct known type (e.g., cell, peak, etc.).

Graph embedding methods learn a *D*-dimensional embedding vector for each *v* ∈ *V* by optimizing a link prediction objective via stochastic gradient descent, with *D*=50 used for our experiments. We will denote the full embedding matrix as *θ* ∈*R*^|*V*|×*D*^ and the embedding for an entity *v* as *θ_v_*.

For an edge *e* = (*u, v*), we denote *s_e_* = *θ_u_* ∗ *θ_v_* as the score for *e*, and optimize a multi-class log loss

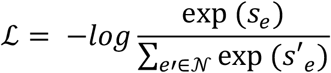

Where 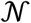 is a set of “negative sampled” candidate edges generated by corrupting *e*^42^. This log loss objective attempts to maximize the score for all (*u, v*) ∈ *E* and minimize it for (*u, v*) ∉ *E*.

Negative samples are constructed by replacing either the source or target entity in the target edge *e* = (*u, v*) with a randomly sampled entity. However, in graphs like ours where only edges between certain entity types are possible, previous work has shown that it is beneficial to optimize the loss only over candidate edges that satisfy the type constraints^43^. Thus, for e.g., a cell-peak edge we only sample negative candidates between cell and peak entities. This modification is crucial in our setting since most randomly selected edges will be of invalid type (e.g., peak-peak), forcing the embeddings to primarily be optimized for irrelevant tasks (e.g., having low dot product between every pair of peaks).

Furthermore, it has been frequently observed that in graphs with wide distribution of node degrees, it is advantageous to sample negatives proportional to some function of the node degree to produce more informative embeddings that don’t merely capture the degree distribution ^13, 44^. For each graph edge in the dataset encountered in a training batch, we produce 100 negatives by corrupting the edge with a source or destination sampled uniformly from the nodes with the correct types for this relation and 100 by corrupting the edge with a source or destination node sampled with probability proportional to its degree^13^.

As with many ML methods, graph embeddings are prone to overfitting in a low-data regime (i.e., low ratio of edges to parameters). We observed overfitting measurable as a gap between training and validation loss on the link prediction task, which we addressed with *L*2 regularization on the embeddings *θ*,

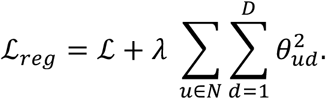

with *λ* = *wd* ∗ *wd_interval*. For weight decay parameter (*wd*), by default it is calculated automatically as 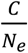, where *N_e_* is the training sample size (i.e., the total number of edges) and *C* is a constant. For weight decay interval (*wd_interval*), we set it to 50 for all experiments.

We use the PyTorch-BigGraph framework, which provides efficient computation of multi-relation graph embeddings over multiple entity types and can scale to graphs with millions or billions of entities^13^. For 1.3 million cells, the PyTorch-BigGraph training itself takes only ~ 1.5 hours using 12 cores without the requirement of GPU (https://simba-bio.readthedocs.io/en/latest/rna_10x_mouse_brain_1p3M.html).

The resulting graph embeddings have two desirable properties that we will take advantage of:

1. First-order similarity: for two entity types *T*_1_, *T*_2_ with a relation between them, edges with high likelihood should have higher dot product; specifically, for any *u* ∈ *T*_1_, the predicted probability distribution over edges to *T*_2_ originating from *u* is approximated as 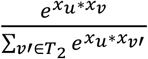.
2. Second-order similarity: within a single entity type, entities that have ‘similar contexts’, i.e., a similar distribution of edge probabilities, should have similar embeddings. Thus, the embeddings of each entity type provide a low-rank latent space that encodes the similarity of those entities’ edge distributions.

### Evaluation of the model during training

During the PyTorch-BigGraph training procedure, a small percent of edges is held out (by default, the evaluation fraction is set to 5%) to monitor overfitting and evaluate the final model. Five metrics are computed on the reserved set of edges, including mean reciprocal rank (MRR, the average of the reciprocal of the ranks of all positives), R1 (the fraction of positives that rank better than all their negatives, i.e., have a rank of 1), R10 (the fraction of positives that rank in the top 10 among their negatives), R50 (the fraction of positives that rank in the top 50 among their negatives), and AUC (Area Under the Curve). By default, we show MRR along with training loss and validation loss while other metrics are also available in SIMBA package (**Supplementary Fig. 1a**). The learning curves for validation loss and these metrics can be used to determine when training has completed. The relative values of training and validation loss along with these evaluation metrics can be used to identify issues with training (underfitting vs overfitting) and tune the hyperparameters weight decay, embedding dimension, and number of training epochs appropriately. For example, in **Supplementary Figure 1a** training can be stopped once the validation loss plateaus. However, for most datasets we find that the default parameters do not need tuning.

### Softmax transformation

PyTorch-BigGraph training provides initial embeddings of all entities (nodes). However, entities of different types (e.g., cells vs peaks, cells of different batches or modalities) have different edge distributions and thus may lie on different manifolds of the latent space. To make the embeddings of entities of different types comparable, we transform the embeddings of features with the Softmax function by utilizing the first-order similarity between cells (reference) and features (query). In the case of batch correction or multi-omics integration, the Softmax transformation is also performed based on the first-order similarity between cells of different batches or modalities.

Given the initial embeddings of cells (reference) (*v*_*c*_1__, …, *v_c_n__*) and features (*V*_*f*_1__, …, *v_d_m__*), the model-estimated probability of an edge (*c_i_*, *f_j_*) obeys

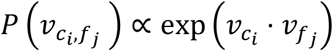

Therefore, if a random edge was sampled from feature *f_j_* to a cell, the model would estimate the distribution over such edges as

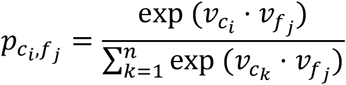

i.e., the Softmax weights between all cells {*c_i_*}and the feature *f_j_*. We can then compute new embeddings for features as a linear combination of the cell embeddings weighted by the edge probabilities raised to some power.

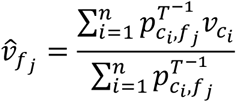

*T* is a temperature hyperparameter that controls the sharpness of the weighting over cells. At *T* = 1, the cell embeddings are weighted by their estimated edge probabilities; at *T* → 0, each feature embedding is assigned the cell embedding of its nearest neighbor; at *T* → ∞, it becomes a discrete uniform distribution, and each query becomes the average of reference embeddings. We set *T* = 0.5 for all the analyses.

These steps are implemented in the function “si.tl.embed()” in the SIMBA package.

### Metrics to assess cell-type specificity

Four metrics are proposed to assess the cell type specificity of each feature from different aspects, including max value (a higher value indicates higher cell-type specificity), Gini index (a higher value indicates higher cell-type specificity), standard deviation (a higher value indicates higher cell-type specificity), and entropy (a lower value indicates higher cell-type specificity). We observe these four metrics generally give consistent results. For SIMBA metric plot, by default, Gini index is plotted against max value. For feature *f_j_*:

The max value is defined as the average normalized similarity of top *k* cells (by default, *k*=50). The similarity normalization function is defined as:

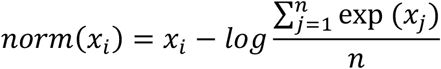

Where *i* = 1, …, *n. n* is the number of cells and *x_i_* represents the dot product of 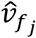 and the embedding of cell *i*.

The max value is computed as:

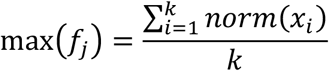

The Gini index is computed as:

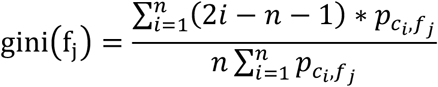

The standard deviation is computed as:

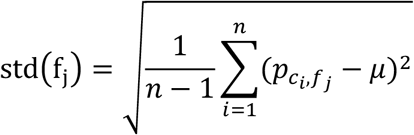

Where 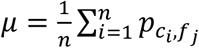.

Entropy is computed as:

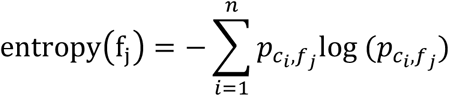

### Queries of entities in SIMBA space

The informative SIMBA embedding space serves as a database of entities including cells and features. To query the “SIMBA database” for the neighboring entities of a given cell or feature, we first build a k-d tree of all entities based on their SIMBA embeddings. We then search for the nearest neighbors in the tree using Euclidean distance. To do so, SIMBA query can perform either K-nearest neighbors (KNN) or nearest neighbor search within a specified radius. SIMBA also provides the option to limit the search to entities of certain types, which is useful when a certain type of entity significantly outnumbered others. For example, the K nearest features of a given cell may be all peaks while genes are the features of interest. In this case, SIMBA allows users to add “filters” to ensure that nearest neighbor search is performed within the specified types of entities. This procedure is implemented in the function “st.tl.query()” and its visualization is implemented in the function “st.pl.query()” in the SIMBA package.

### Identification of master regulators

To identify master regulators, we take into consideration both the cell type specificity of each pair of TF motif and TF gene and the distance between them. More specifically, for each TF motif, first its distances (Euclidean distance by default) to all the genes are calculated in the SIMBA embedding space. Then the rank of this TF gene among all these genes is computed. In addition, we also assess the cell type specificity of this pair of TF motif and TF gene based on SIMBA metrics (by default, max value and Gini index are used). The same procedure is performed for all TFs. Finally, we identify master regulators by filtering out TFs with low cell-type specificity and scoring them based on TF gene rank. This procedure is implemented in the function “st.tl.find_master_regulators()” in SIMBA package.

### Identification of TF target genes

Given a master regulator, its target genes are identified by comparing the locations of the TF gene, TF motif, and the peaks near the genomic loci of candidate target genes in the SIMBA co-embedding space (**Fig. 4e**). More specifically we first search for *k* nearest neighbor genes around the motif (TF motif) and the gene (TF gene) of this master regulator, respectively (*k* = 200 by default). The union of these neighbor genes is the initial set of candidate target genes. These genes are then filtered based on the criterion that open regions (peaks) within 100kb upstream and downstream of the TSS of a putative target gene must contain the TF motif.

Next, for each candidate target gene, we compute four types of distances in SIMBA embedding space: distances between the embeddings of 1) the candidate target gene and TF gene; 2) the candidate target gene and TF motif; 3) peaks near the genomic locus of the candidate target gene and TF motif; 4) peaks near the genomic locus of the candidate target gene and the candidate gene. All the distances (Euclidean distances by default) are converted to ranks out of all genes or all peaks to make the distances comparable across different master regulators.

The final list of target genes is decided using the calculated ranks based on two criteria: 1) at least one of the nearest peaks to TF gene or TF motif is within a predetermined range (top 1,000 by default); 2) the average rank of the candidate target gene is within a predetermined range (top 5,000 by default). This procedure is implemented in the function “st.tl. find_target_genes ()” in SIMBA.

### Benchmarking scATAC-seq computational methods

To compare SIMBA to other scATAC-seq computational methods including SnapATAC ^4^, Cusanovich2018^45^, and cisTopic^46^, we employed the previously developed benchmarking framework from Chen et al^14^ (**Supplementary Table 1**). This framework evaluates different methods based on their ability to distinguish cell types. We applied three clustering algorithms: k-means clustering, hierarchical clustering, and Louvain on the feature matrix derived from each method.

For datasets with ground-truth (FACS-sorted labels or known tissue labels), including simulated bone marrow data, Buenrostro 2018, and sci-ATAC-seq subset, three metrics including Adjusted Rand Index (ARI), Adjusted Mutual Information (AMI), and Homogeneity are applied to evaluate the performance. ARI measures the similarity between two clusters, comparing all pairs of samples assigned to matching or different clusters in the predicted clustering solution vs the true cluster/cell type label. AMI describes an observed frequency of co-occurrence compared to an expected frequency of co-occurrence between two variables, informing the mutual dependence or strength of association of these two variables. Homogeneity measures whether a clustering algorithm preserves cluster assignments towards samples that belong to a single class. A higher metric value indicates a better clustering solution.

For 10x PBMCs dataset with no ground truth, the Residual Average Gini Index (RAGI) proposed in the benchmarking study^14^ is used as the clustering evaluation metric. RAGI measures the relative exclusivity of marker genes to their corresponding clusters in comparison to housekeeping genes, which should demonstrate low specificity to any given cluster. In brief, the mean Gini Index is computed for both marker genes and housekeeping genes. The difference between the means is computed to obtain the average residual specificity (i.e., RAGI) of a clustering solution with respect to marker genes. A higher RAGI indicates a better separation of biologically distinct clusters.

### Benchmarking single-cell batch correction methods

The batch correction performance of SIMBA was compared to Seurat3^12^, LIGER^11^ and Harmony^10^ in two benchmark datasets: the mouse atlas dataset and the human pancreas dataset (**Supplementary Table 1**). For Seurat3, LIGER and Harmony, the batch correction was done with the same parameters used in a previous benchmark study^25^.

To evaluate the batch integration performance, average Silhouette width (ASW), adjusted Rand index (ARI), and local inverse Simpson’s index (LISI)^10^ were calculated for the batches and cell types using the Euclidean distance as described in a previous benchmark^25^. To make a fair evaluation, only the cell types that are present in all batches were considered. We used the same number of dimensions (50) for these methods and all other parameters were set as in the benchmark.

#### Average Silhouette width (ASW)

Average Silhouette width is the mean value of Silhouette scores calculated from each cell. Silhouette width measures the relative closeness of cells with the same label compared to the cells with the different label and ranges from −1 to +1. Silhouette score for a data point with a label is calculated as

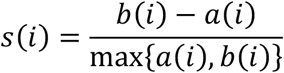

where *a*(*i*) is the distance to the closest point with the same label, and *b*(*i*) is the distance to the closest point with different labels. A high Silhouette score means the point is located more closely with the same label, where a low Silhouette score closer to −1 means the point is located closer with different labels than that of itself. The ideal batch correction result will give a low ASW score for batch labels as the point is well mixed with other batches and a high ASW score for the cell type labels as the cells of the same cell type should cluster together after the batch correction. The final score is calculated as the median ASW scores from 20 subsets of randomly sampled 80% cells.

#### Average Rand Index (ARI)

To evaluate the cell type purity, the true cell type labels and the k-means clustering solution were used to calculate the cell type ARI. To evaluate the batch correction performance, the true batch labels and the k-means clustering solution were used to calculate the batch ARI. The final ARI was calculated as the median ARI scores of 20 subsets comprised of randomly sampled 80% cells for batches and cell types, respectively. A superior batch correction will have a high cell type ARI (high agreement between the clustering solution and the true cell type labels), and a low batch ARI (the clustering solution is not mainly driven by batches and clusters contain cells with well-mixed batch labels).

#### Local Inverse Simpson’s Index (LISI)

Local Inverse Simpson’s Index (LISI) ^10^ measures the local batch and cell type mixing. For each data point, it considers the Gaussian kernel weighted distribution of labels in its neighborhood with a perplexity argument. We set perplexity to 50 40 as in the previous benchmark study. Using the weighted neighborhood label distribution, the inverse Simpson’s index is calculated as 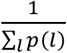 where *l* is the batch or cell type labels and *p*(*l*) is the probability of each label in the local neighborhood obtained with the kernel. For each cell, the LISI is the expected number of cells to be sampled locally before a cell of the same label is sampled. A perfect batch correction will have a cell type LISI (cLISI) of 1 and a batch LISI (integration LISI, iLISI) close to the number of batches. The final LISI score was calculated as the average LISI scores of all cells.

Further details are described in **Supplementary Note 2**.

### Benchmarking single cell multi-omics integration methods

Two pairs of scRNA-seq and scATAC-seq datasets manually split from the dual-omics SHARE-seq mouse skin dataset and 10X PBMCs dataset respectively were used for the modality integration task. For Seurat3 and LIGER, the parameters and preprocessing were done as described in their documentations. However, for the LIGER analysis of the SHARE-seq mouse skin dataset the parameter ‘lambda’ was set to 30 and the ‘ref_dataset’ was set to scATAC-seq to get a better alignment. For the Raw results, the activity matrix of scATAC-seq was constructed using Seurat3 and the first 20 PCs of the scRNA-seq count matrix and the activity matrix were used for the comparison. The integration results generated by each method were evaluated with four metrics— Anchoring distance, anchoring distance rank, Silhouette index, and cluster agreement— as described below.

#### Anchoring distance

The Anchoring distance was proposed in Dou et al., 2020^47^ and is the normalized distance between the matched cells of two modalities (e.g. RNA and ATAC). Here we considered the Euclidean distance and normalized the distance by the mean of the distances calculated between random pairs of cells. The number of pairs randomly sampled was set to 10% of the total number of cells.

#### Anchoring distance rank

Given that the anchoring distance does not account for the local density of cells, we propose a new metric entitled *anchoring distance rank* (ADR). The ADR is based on the normalized rank of the distance between the matched cells of two modalities. For each cell *x_ij_* with cell identity i and modality j, the distance between the cell and all the other cells of the other modality j’, *d*(*x_ij_*, *x_kj′_*),*k* = 1,…, *N* is calculated, where N is the total number of cells. Then the rank of *r_i_* = *d*(*x_ij_*, *x_ij′_*) within the calculated distances is normalized by the number of pairs *N* − 1 to obtain the final anchoring rank 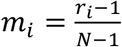. For each cell, an anchoring rank of 0 indicates an ideal modality integration performance as the matched cells are closest to each other in the embedding.

#### Silhouette index

The silhouette index was calculated as described in 10) based on the cluster assignment wherein each cluster consists of two cells, one cell from a scRNA-seq dataset and one cell from a scATAC-seq dataset.

#### Fraction in the same cluster

Fraction in the same cluster was calculated as the fraction of the matched cells from two modalities in the same cluster. The clusters of cells were generated using Louvain algorithm and the number of clusters is equal to the number of cell types in the dataset.

Further details are described in **Supplementary Note 3**.

## Data availability

All the datasets used in this study (eight scRNA-seq datasets, four scATAC-seq datasets, and three dual-omics datasets) are summarized in **Supplementary Table 1**. All these datasets are curated in the SIMBA package, and they can be easily downloaded and imported directly to reproduce the analyses presented in this manuscript.

## Code availability

We provide a comprehensive Python package ‘simba’ available at https://anaconda.org/bioconda/simba and https://github.com/pinellolab/simba. All the proposed procedures are implemented in the “simba” package. ‘simba’ can be easily installed with conda “*conda install simba*”. We also built a website (https://simba-bio.readthedocs.io), providing a detailed introduction of the ‘simba’ software and several SIMBA tutorials for different types of single-cell analyses presented in this manuscript.

## Acknowledgements

The authors thank Ledell (Yu) Wu, Facebook, for the helpful discussions about Starspace; Dr.Sai Ma and Dr. Jason Buenrostro, Broad Institute of MIT and Harvard, for sharing the SHARE-seq data and metadata. The authors also acknowledge members of the Pinello lab, Mass General Hospital/Harvard Medical School, for helpful comments and feedback. This project has been made possible in part by grant number 2019-202669 from the Chan Zuckerberg Foundation to LP. LP is also partially supported by the National Human Genome Research Institute (NHGRI) Genomic Innovator Award (R35HG010717). M.E.V. is supported by the National Institutes of Health (NIH) under the Ruth L. Kirschstein National Research Service Award (1F31CA257625-01) from the National Cancer Institute (NCI).

## Author contributions

H.C. and L.P. conceived this project. H.C. developed SIMBA, wrote the SIMBA package, and performed SIMBA analysis on all datasets. A.L. contributed to the adaption of PyTorch-BigGraph to single cell analysis. J.R. and H.C. performed the comparison analysis on batch correction and data integration. M.E.V. and H.C. performed the comparison analysis on scATAC-seq data. L.P. and A.L. provided guidance and supervised this project. All the authors wrote and approved the final manuscript.

## Supplementary Figures

**Supplementary Figure 1.**
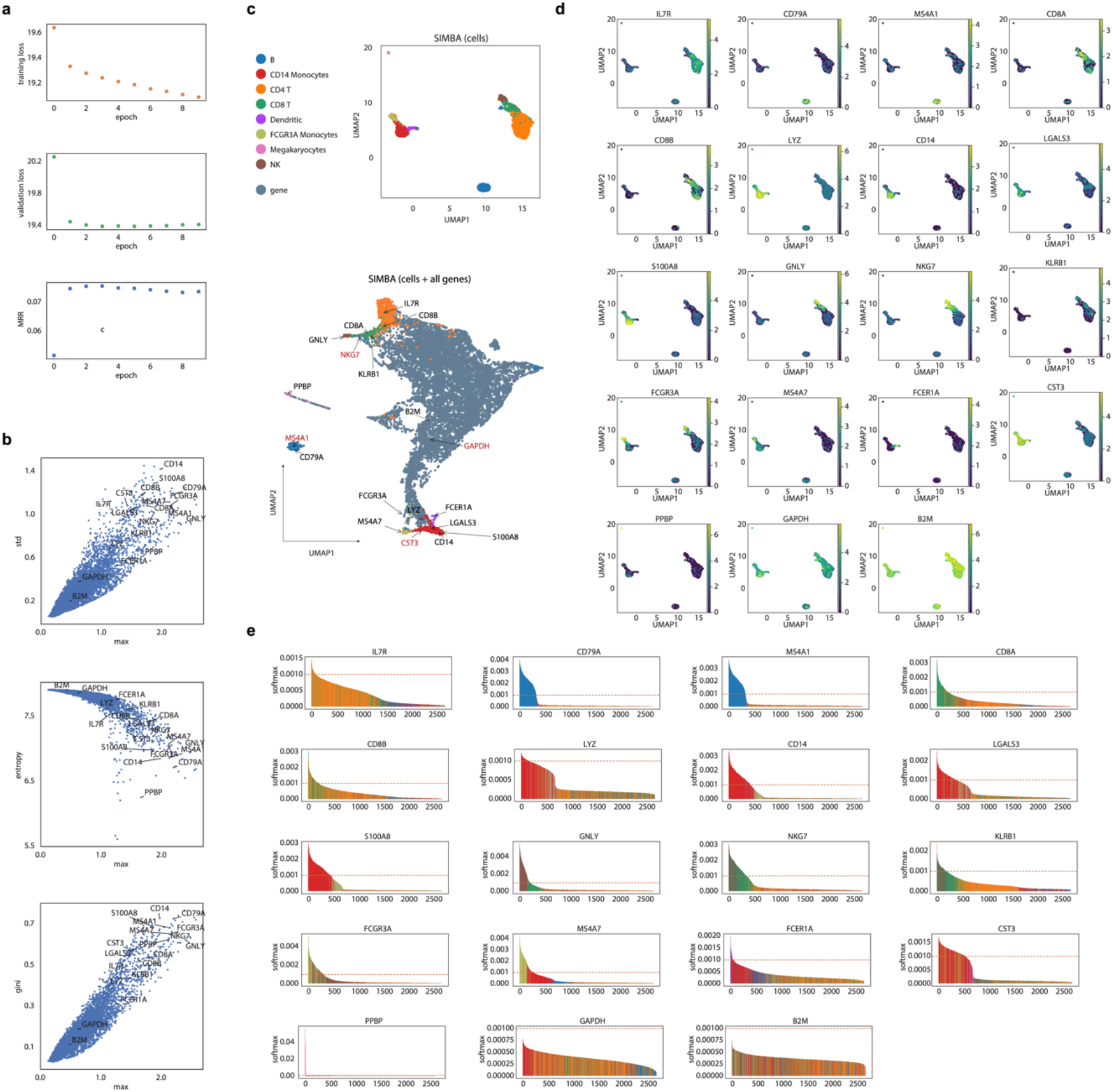
SIMBA analysis of the scRNA-seq 10x PBMCs dataset. **a.** Three default metrics used to evaluate SIMBA training procedure, including training loss (top), validation loss (middle), mean reciprocal rank (MRR) **b.** SIMBA metric plots of genes. All the genes are plotted according to the Gini index against max score, standard deviation (std) against max score, and entropy against max score, respectively. The same set of genes as in Figure 2c are highlighted. **c.** UMAP visualization of the SIMBA embeddings of cells and the SIMBA embeddings of cells and all genes. Genes highlighted in (b) are also highlighted in the UMAP plot. **d.** UMAP visualization of the SIMBA embeddings of cells, colored by gene expression of the genes highlighted in (b). **e.** SIMBA barcode plots of the genes highlighted in (b).

**Supplementary Figure 2.**
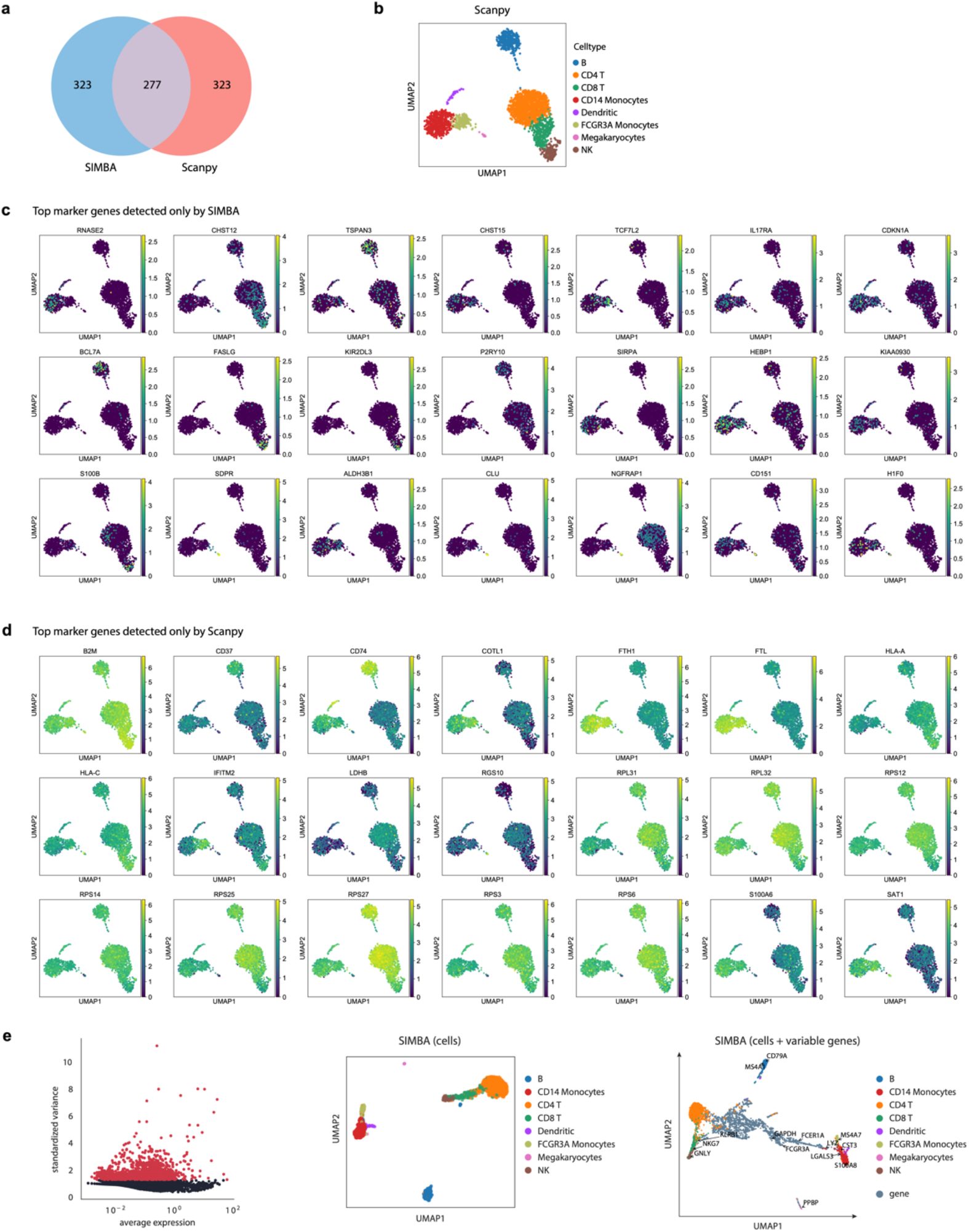
Comparison of SIMBA with Scanpy on the scRNA-seq 10x PBMCs dataset. **a.** Venn diagram of top marker genes identified by SIMBA and Scanpy **b.** Scanpy-derived UMAP visualization of cells colored by cell type **c.** Top marker genes detected only by SIMBA. Colored by intensity of gene expression. **d.** Top marker genes detected only by Scanpy. Colored by intensity of gene expression. **e.** SIMBA embedding result after implementing variable gene selection. Left: variable gene selection step implemented in SIMBA. Middle: UMAP visualization of SIMBA embeddings of cells. Right: UMAP visualization of SIMBA embeddings of cells and variable genes.

**Supplementary Figure 3.**
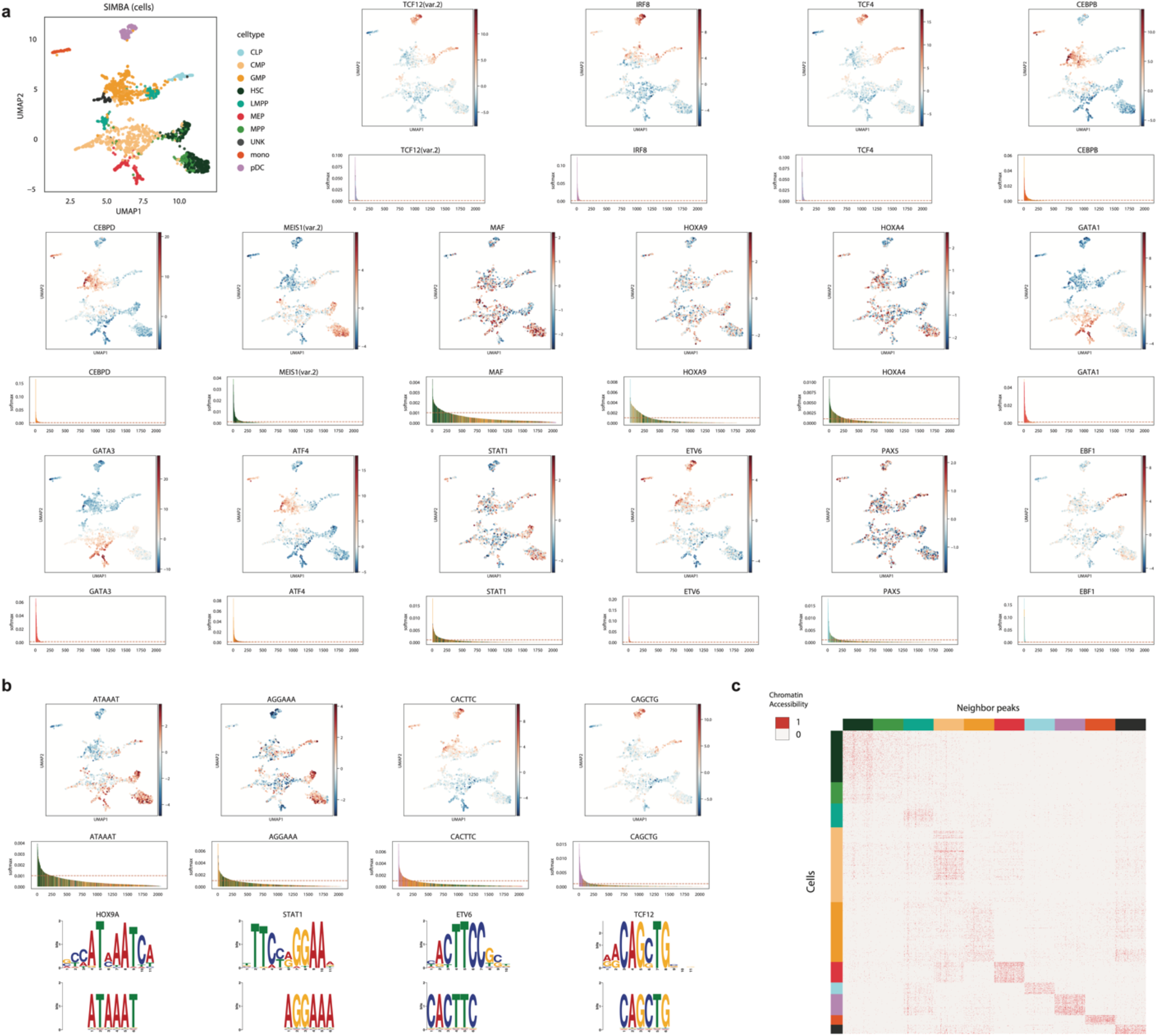
SIMBA analysis of the *Buenrostro2018* dataset **a.** UMAP visualization of SIMBA embeddings of cells colored by cell type (top-left), and TF activity scores of TF motifs calculated with chromVAR, respectively. The SIMBA barcode plot of each TF motif is shown below the UMAP plot. **b.** Top: UMAP visualization of SIMBA embeddings of cells colored by TF activity scores of k-mers calculated with chromVAR. Middle: SIMBA barcode plots of the corresponding k-mers. Bottom: the matching known motif against the enriched k-mer sequences. **c.** Heatmap of cells against neighboring peaks of each cell type that are selected in the SIMBA co-embedding space. Chromatin accessibility is binary and colored accordingly.

**Supplementary Figure 4.**
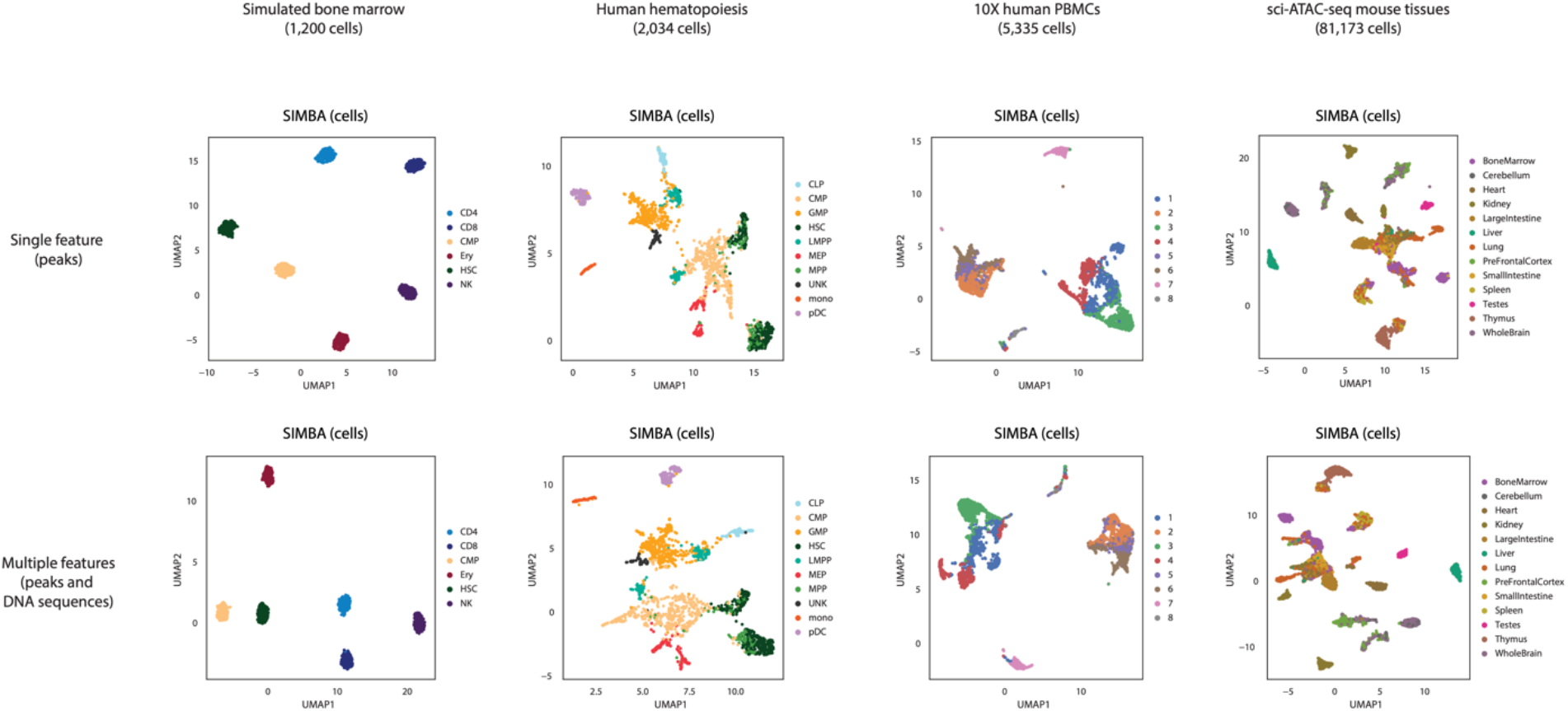
Comparison of SIMBA performance using scATAC-seq peaks and DNA sequence content vs only scATAC-seq peaks. **Top**: UMAP visualization of SIMBA embeddings of cells for each indicated dataset generated from only scATAC-seq peak information. **Bottom**: UMAP visualization of SIMBA embeddings of cells for each indicated dataset generated from scATAC-seq peak information and DNA sequence content information.

**Supplementary Figure 5.**
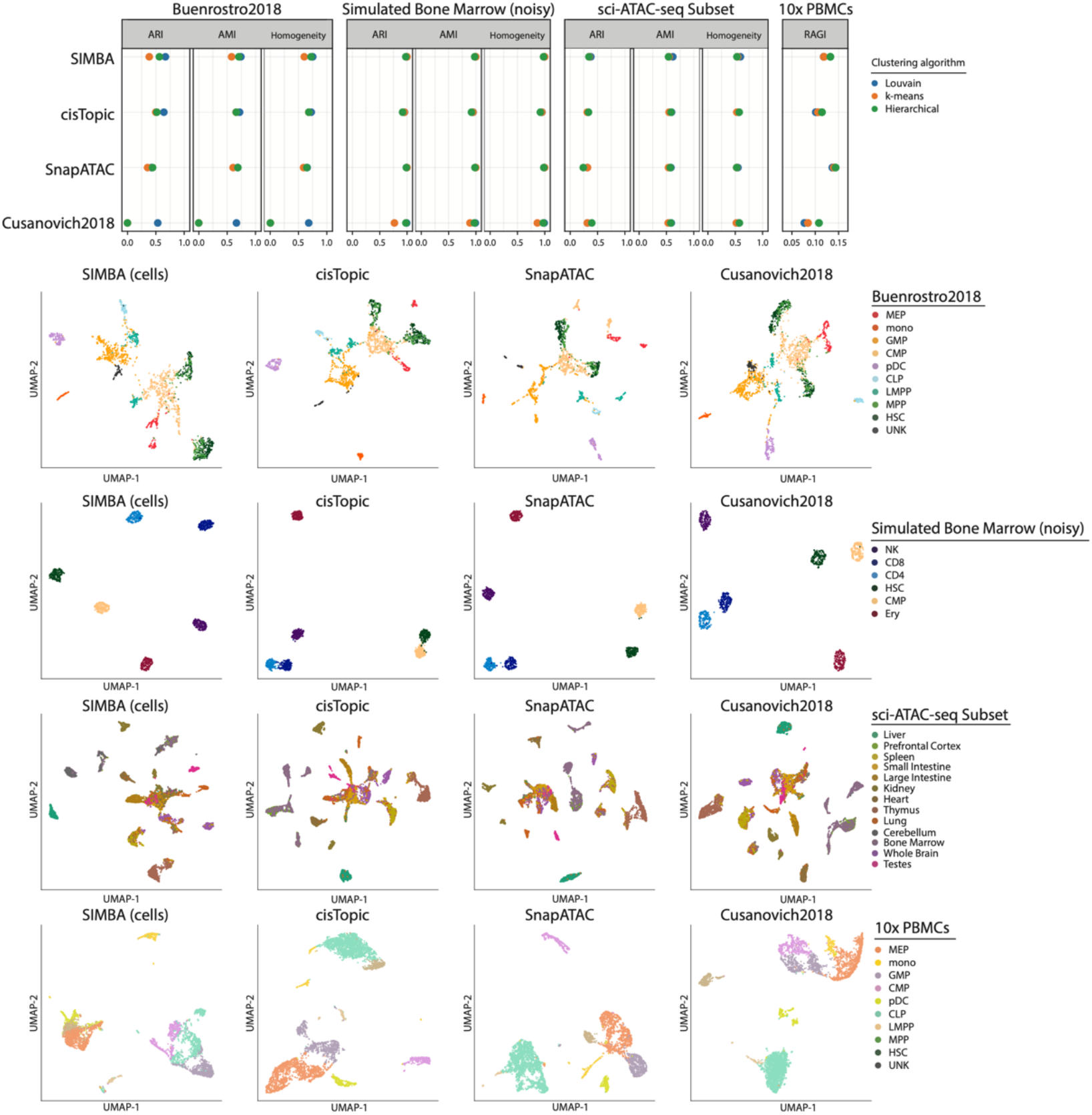
Benchmark of SIMBA against top-performing scATAC-seq analysis methods. Top: Evaluation of SIMBA and other methods including cisTopic, SnapATAC, *Cusanovich2018* for scATAC-seq analysis using metrics 1) ARI, AMI, and Homogeneity for datasets with ground truth cell type labels and 2) Residual Average Gini Index (RAGI) for the 10x PBMCs dataset without ground truth labels. Bottom: UMAP visualization of feature matrices produced by each method on each dataset colored by cell type annotation or cluster label.

**Supplementary Figure 6.**
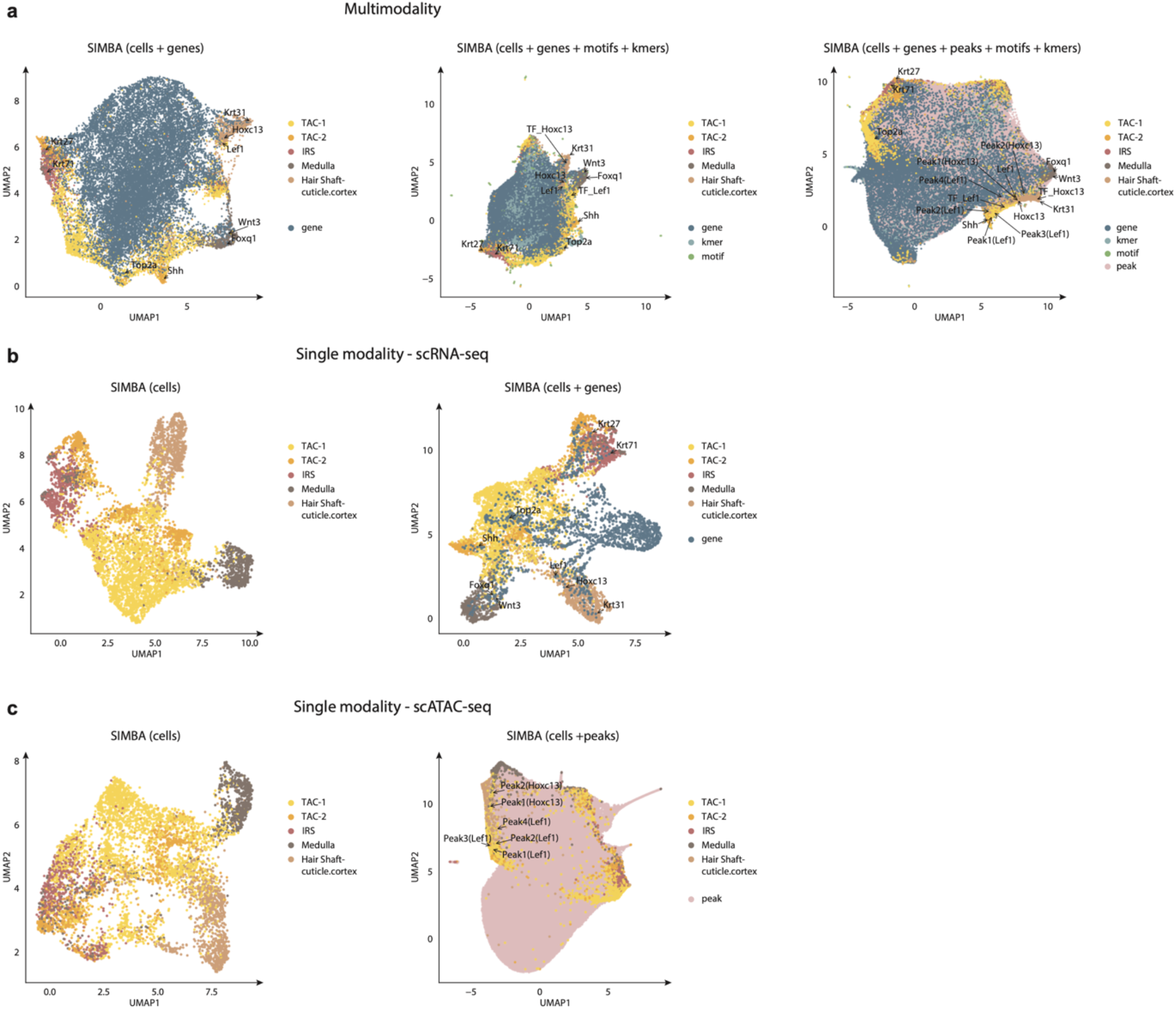
SIMBA multimodal analysis of the SHARE-seq hair follicle dataset. **a.** SIMBA embedding results when both gene expression and chromatin accessibility are encoded in the graph. Left: UMAP visualization of SIMBA embeddings of cells and genes. Middle: UMAP visualization of SIMBA embeddings of cells along with genes, TF motifs, and k-mers. Right: UMAP visualization of SIMBA embeddings of cells along with genes, peaks, TF motifs, and k-mers. **b.** SIMBA embedding results when only gene expression is encoded in the graph. Left: UMAP visualization of SIMBA embeddings of cells. Right: UMAP visualization of SIMBA embeddings of cells and variable genes. **c.** SIMBA embedding results when only chromatin accessibility is encoded in the graph. Left: UMAP visualization of SIMBA embeddings of cells. Right: UMAP visualization of SIMBA embeddings of cells and peaks.

**Supplementary Figure 7.**
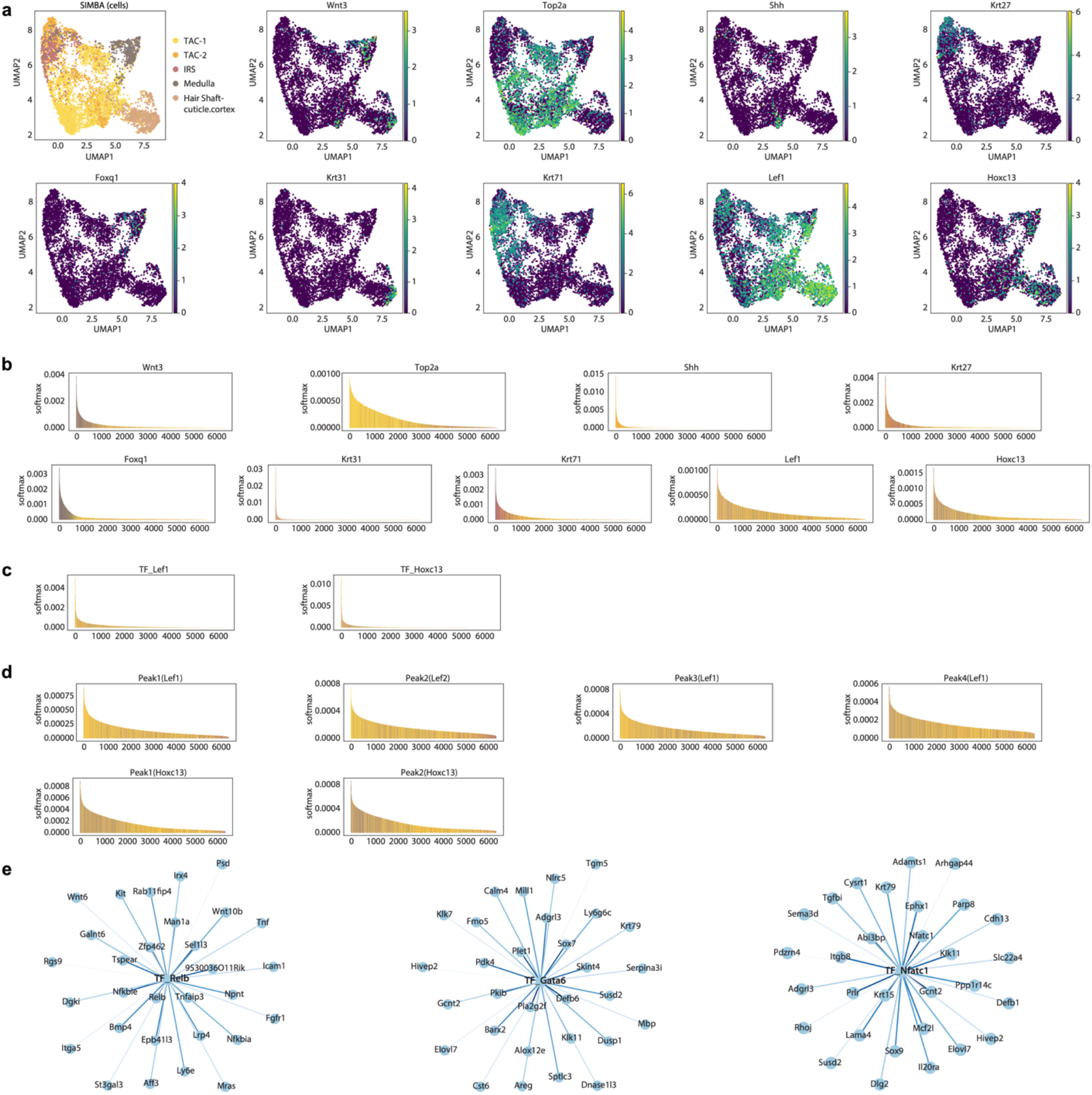
Cell type specific marker genes and the target genes of master regulators identified by SIMBA in the SHARE-seq hair follicle subset dataset. **a.** UMAP visualization of SIMBA embeddings of cells colored by cell type and gene expression intensity. **b.** SIMBA barcode plots of each gene plotted above. **c.** SIMBA barcode plots of TF motifs *Lef1* and *Hoxc13*. **d.** SIMBA barcode plots of peaks near the loci of *Lef1* and *Hoxc13*. **e.** Top 30 target genes of the master regulators *Relb, Gata6*, and *Nfatc1* as inferred by SIMBA.

**Supplementary Figure 8.**
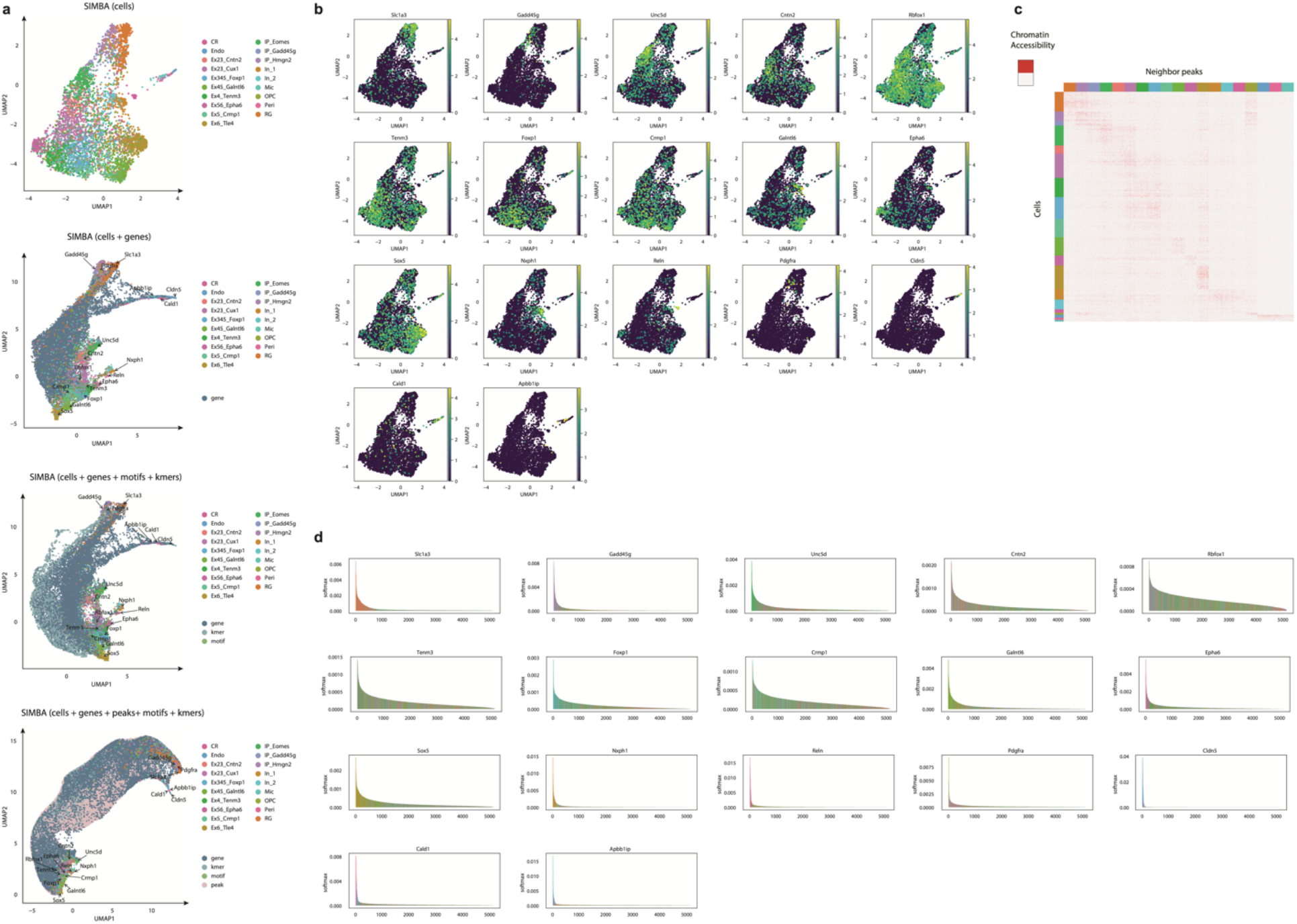
SIMBA multimodal analysis of the SNARE-seq mouse cerebral cortex dataset. **a.** From top to bottom: UMAP visualization of SIMBA embeddings of (i) cells (ii) genes alongside cells (iii) genes, motifs, and k-mers alongside cells (iv) genes, peaks, motifs, and k-mers alongside cells. **b.** UMAP visualization of SIMBA embeddings of cells colored by indicated gene expression intensity. **c.** Heatmap of cells against neighboring peaks of each cell type that are selected in the SIMBA co-embedding space. Chromatin accessibility is binary and colored accordingly. **d.** SIMBA barcode plots of the genes highlighted in (a).

**Supplementary Figure 9.**
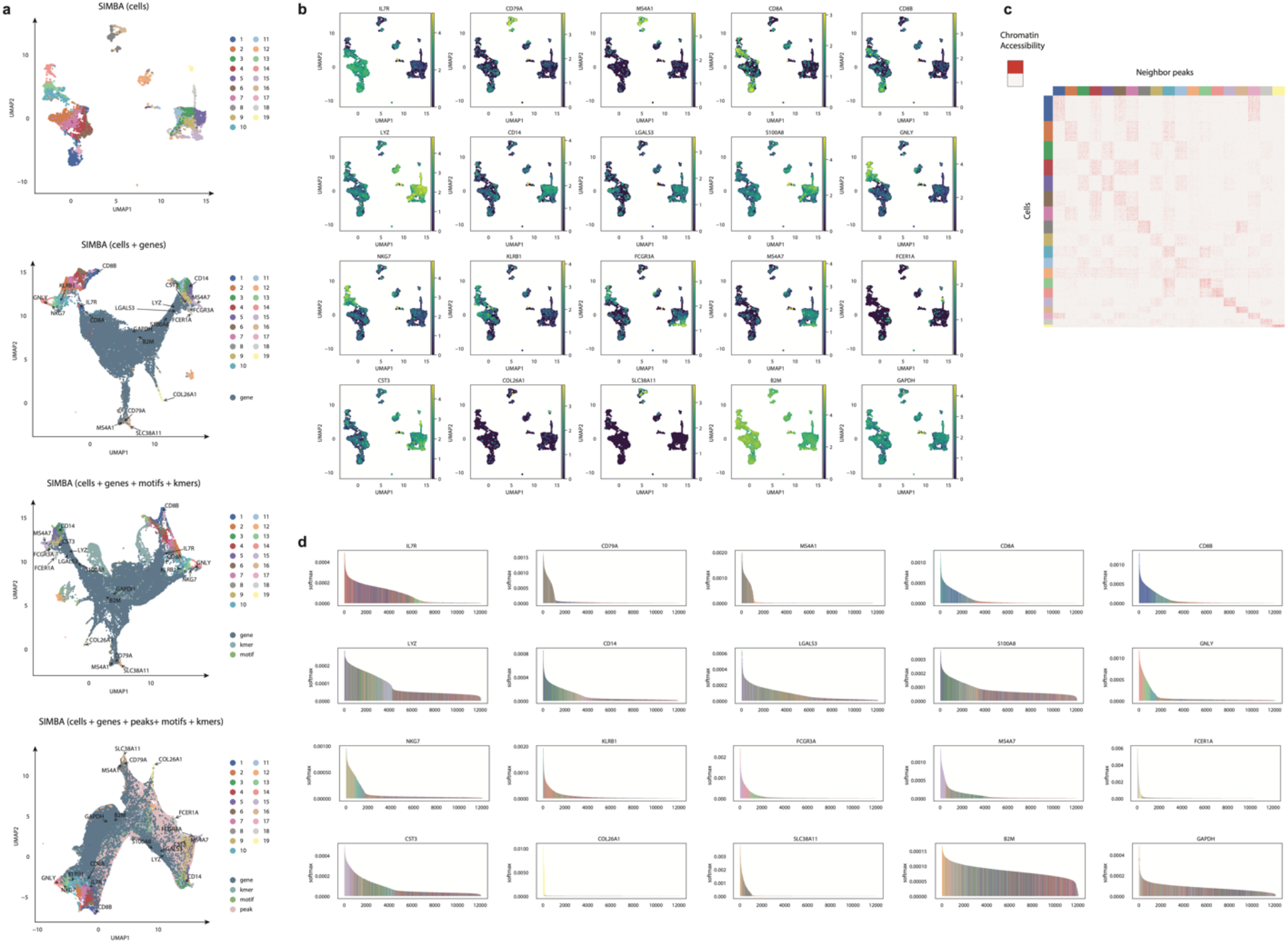
SIMBA multimodal analysis of the 10x multiome PBMCs dataset. **a.** From top to bottom: UMAP visualization of SIMBA embeddings of (i) cells (ii) genes alongside cells (iii) genes, motifs, and k-mers alongside cells (iv) genes, peaks, motifs, and k-mers alongside cells. **b.** UMAP visualization of SIMBA embeddings of cells colored by indicated gene expression intensity. **c.** Heatmap of cells against neighboring peaks of each cluster that are selected in the SIMBA co-embedding space. Chromatin accessibility is binary and colored accordingly. **d.** SIMBA barcode plots of the genes highlighted in (a).

**Supplementary Figure 10.**
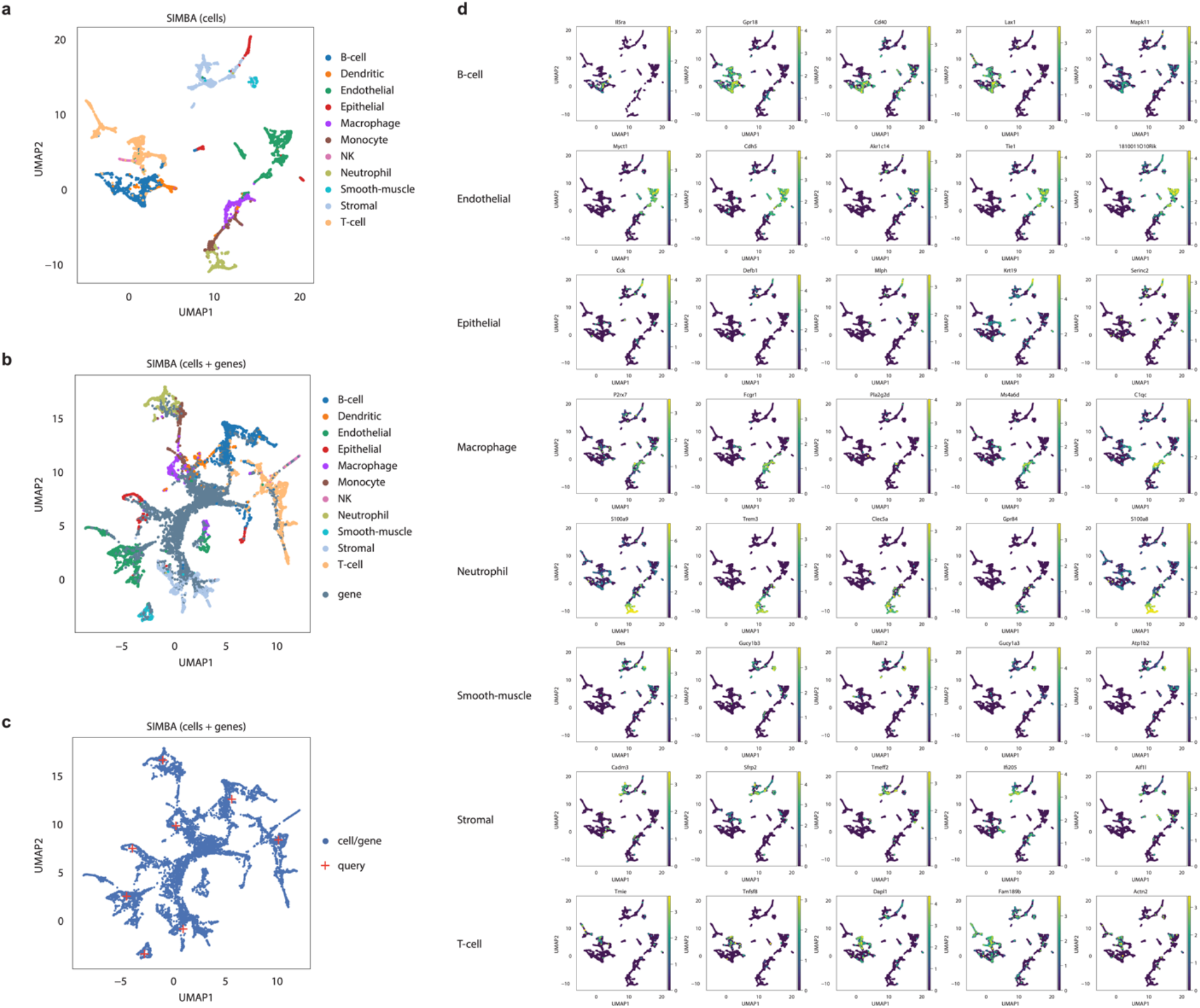
SIMBA-inferred marker genes for the scRNA-seq mouse atlas dataset in batch correction analysis. **a.** UMAP visualization of SIMBA embeddings of cells colored by cell type. **b.** UMAP visualization of SIMBA embeddings of cells and genes. **c.** UMAP visualization of SIMBA embeddings of cells and genes. Biological “query” points are highlighted with a red Nearby informative genes are colored accordingly. **d.** UMAP visualization of SIMBA embeddings of cells colored by indicated gene expression intensity, separated by cell type.

**Supplementary Figure 11.**
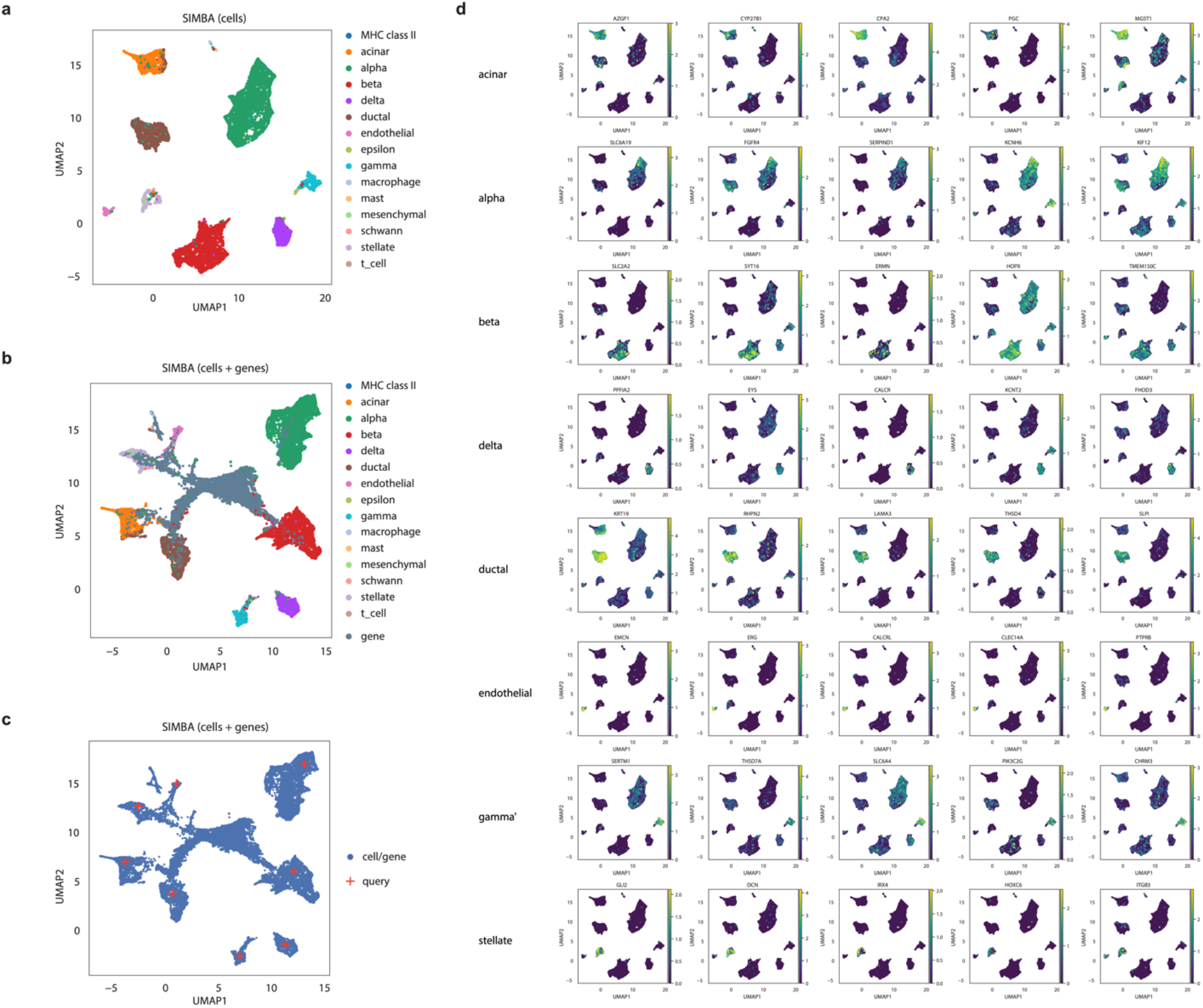
SIMBA-inferred marker genes for the scRNA-seq human pancreas dataset in batch correction analysis. **a.** UMAP visualization of SIMBA embeddings of cells colored by cell type. **b.** UMAP visualization of SIMBA embeddings of cells and genes. **c.** UMAP visualization of SIMBA embeddings of cells and genes. Biological “query” points are highlighted with a red “+”. Nearby informative genes are colored accordingly. **d.** UMAP visualization of SIMBA embeddings of cells colored by indicated gene expression intensity, separated by cell type.

**Supplementary Figure 12.**
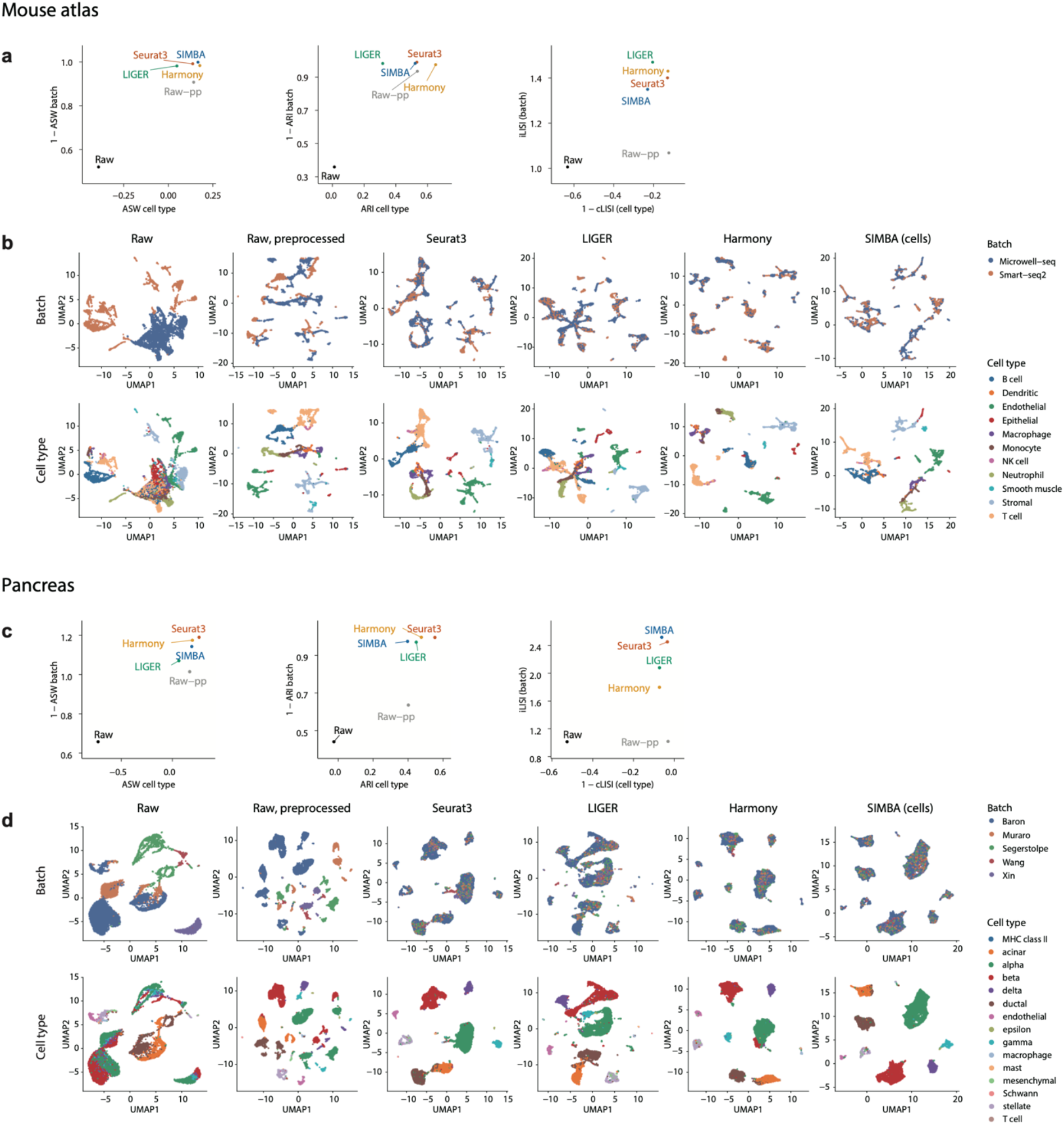
Comparison of SIMBA to other methods for batch correction of the mouse atlas (a-b) and human pancreas scRNA-seq datasets (c-d). **a, c**. Quantitative comparison of SIMBA with three other batch correction methods including Seurat3, LIGER and Harmony, using, left-to-right: average silhouette width (ASW), adjusted Rand index (ARI), and local inverse Simpson’s index (LISI) **b, d**. UMAP visualization of raw and preprocessed data alongside the batch corrected results produced by Seurat3, LIGER, Harmony, and SIMBA. Colored by technology (top) and cell type (bottom).

**Supplementary Figure 13.**
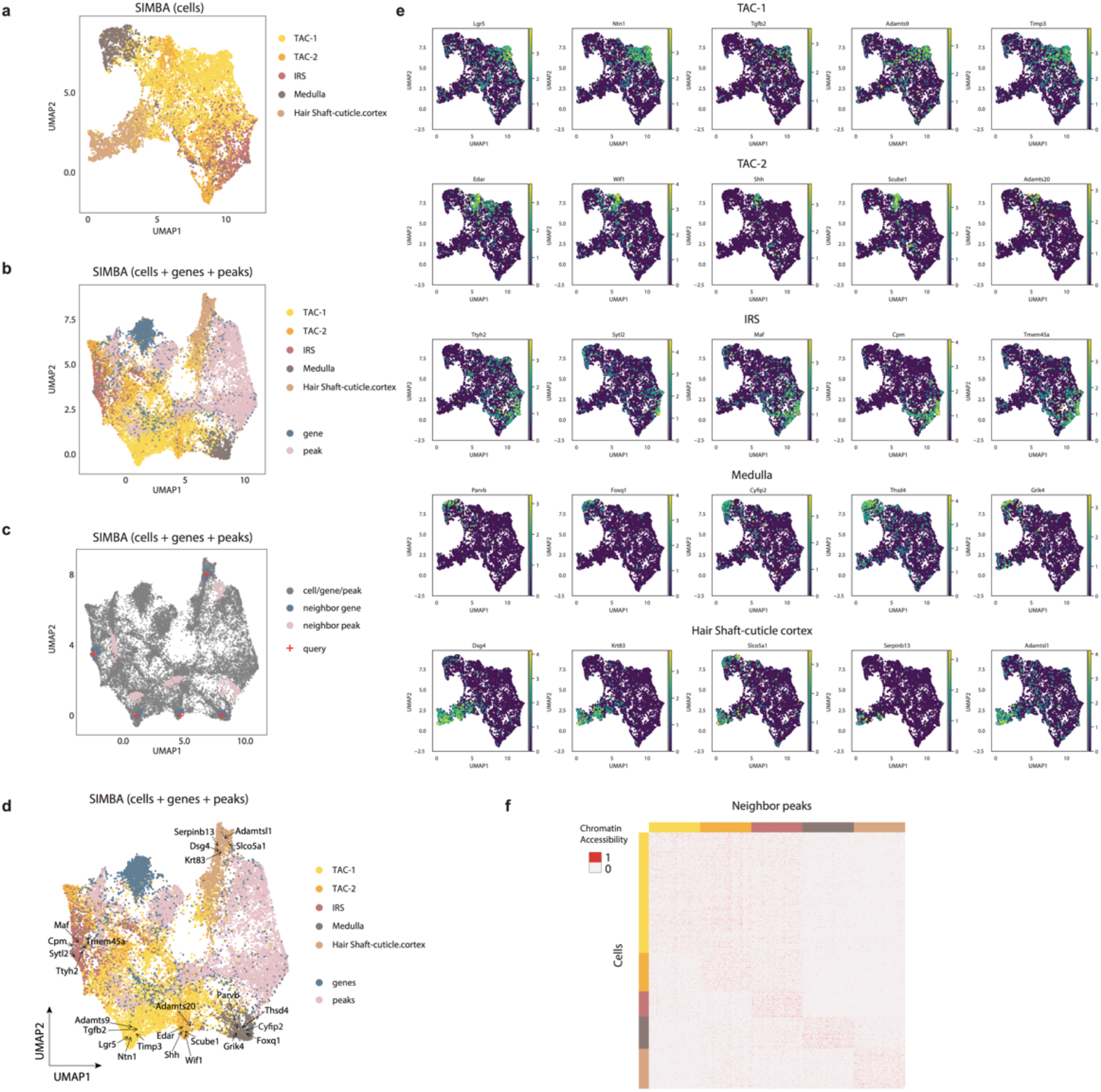
SIMBA-inferred marker features for the SHARE-seq mouse skin dataset in multi-omics integration analysis. **a.** UMAP visualization of SIMBA embeddings of cells with two cellular modalities integrated. **b.** UMAP visualization of SIMBA embeddings of cells, genes, and peaks with two cellular modalities integrated. **c.** UMAP visualization of SIMBA embeddings of cells, genes, and peaks with two cellular modalities integrated. Biological “query” points are highlighted with a red “+”. Nearby informative genes and peaks are colored accordingly. **d.** UMAP visualization of SIMBA embeddings of cells, genes, and peaks with two cell modalities integrated and known marker genes highlighted. **e.** UMAP visualization of SIMBA embeddings of cells colored by indicated gene expression intensity, separated by cell type. **f.** Heatmap of cells against neighboring peaks of each cell type that are selected in the SIMBA co-embedding space. Chromatin accessibility is binary and colored accordingly.

**Supplementary Figure 14.**
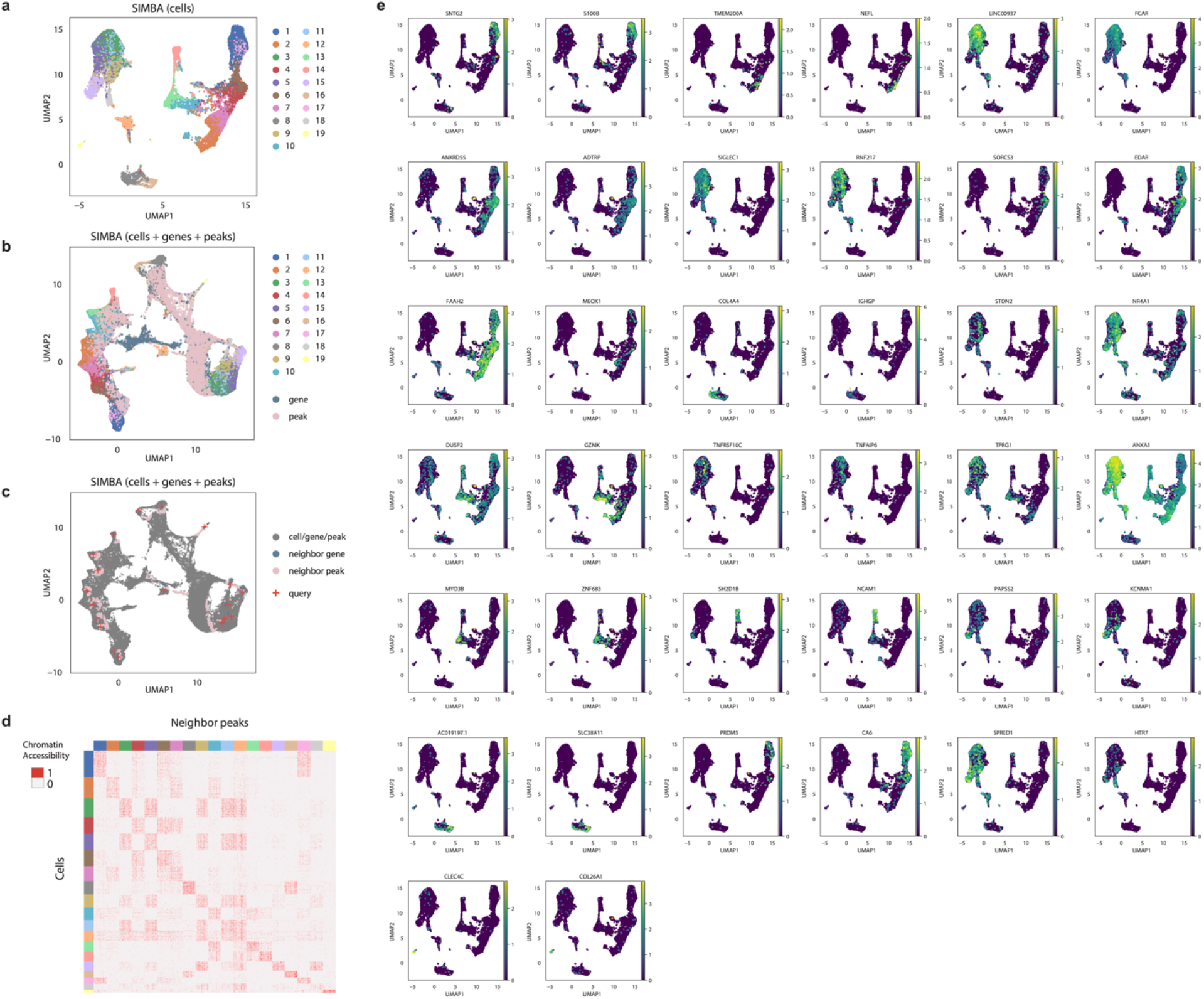
SIMBA-inferred marker features for the 10x human PBMCs dataset in multi-omics integration analysis. **a.** UMAP visualization of SIMBA embeddings of cells with two cellular modalities integrated. **b.** UMAP visualization of SIMBA embeddings of cells, genes, and peaks with two cellular modalities integrated. **c.** UMAP visualization of SIMBA embeddings of cells, genes, and peaks with two cellular modalities integrated. Biological “query” points are highlighted with a red “+”. Nearby informative genes and peaks are colored accordingly. **d.** Heatmap of cells against neighboring peaks of each cluster that are selected in the SIMBA co-embedding space. Chromatin accessibility is binary and colored accordingly. **e.** UMAP visualization of SIMBA embeddings of cells colored by indicated gene expression intensity, separated by cell type.

**Supplementary Figure 15.**
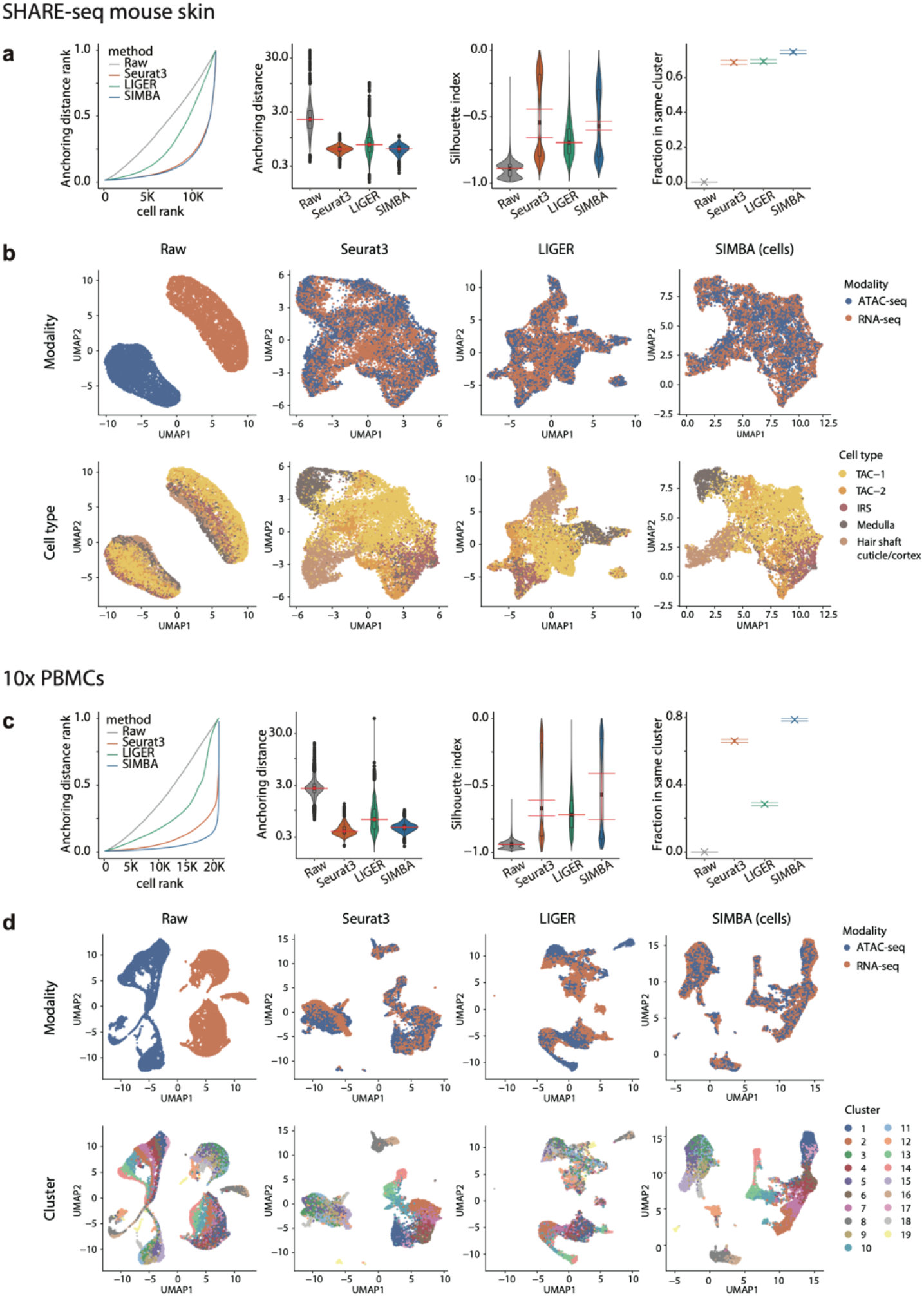
Comparison of SIMBA to other methods for multi-omics integration of the SHARE-seq mouse skin (a-b) and 10x multiome human PBMCs (c-d) datasets. **a, c**. Quantitative comparison of SIMBA with two other methods including Seurat3, LIGER for multi-omics integration, using, left-to-right: anchoring distance rank, anchoring distance, silhouette index, and Fraction in the same cluster. **b, d**. UMAP visualization of the raw scRNA-seq and scATAC-seq data from the 10x multiome human PBMCs dataset alongside the integrated results produced by Seurat3, LIGER, and SIMBA. Colored by data modality (top) and cluster assignment (bottom). The red intervals of violin plot of Anchoring distance and Silhouette index shows the 95% of the mean.

## Supplementary Notes

### Supplementary Note 1: Comparison with scATAC-seq methods

To assess SIMBA’s ability to cluster cell types based on scATAC-seq profiles, we compared SIMBA with specialized methods specifically designed for this task. We observed that SIMBA yields consistent embeddings of cells when using either a single feature (peaks) or multiple features (peaks and DNA sequences from within those peaks) as input to the graph. This comparison was performed across four scATAC-seq datasets of varying profiling technologies and organisms (**Supplementary Fig. 4**). Given these differences, to create a fair comparison we used the same set of features (i.e., peaks) for SIMBA as other methods. SIMBA’s performance was compared against three of the top methods, including SnapATAC^1^, *Cusanovich2018*^2^, and cisTopic^3^ recommended by our recent benchmark study^4^. This comparison was first made qualitatively based on UMAP visualization and then quantitatively based on clustering performance. SIMBA performed as well as or better than each of the methods evaluated. These results comparing SIMBA to scATAC-seq-specialized methods highlight SIMBA’s wide utility for single-cell analyses (**Supplementary Fig. 5**).

### Supplementary Note 2: Comparison with batch correction methods

Multiple methods have now been developed to correct for the technical effects of sample preparation and data collection in single cells. To assess SIMBA’s performance in removing batch effects, we compared it to Seurat3^5^, LIGER^6^ and Harmony^7^, three top-performing batch correction methods recommended in a recent benchmark study^8^.

Two datasets, including a mouse atlas dataset and a human pancreas dataset (see **Supplementary Table 1**), were used for the evaluation. The mouse atlas dataset is composed of two scRNA-seq subsets with shared cell types from different sequencing platforms. The human pancreas dataset is composed of five samples pooled from five distinct sources using four different sequencing techniques wherein not all cell types are shared across each sample.

To qualitatively compare these methods, we visualized cells of each dataset before and after batch-correction in UMAP plots (**Supplementary Fig. 12b,d**). To quantitatively evaluate the performance of each method, using the benchmarking pipeline laid out in Tran *et al*^8^, we measured the conservation of biological information and batch effect removal based on three different metrics: average silhouette width (ASW), adjusted Rand index (ARI), and local inverse Simpson’s index (LSI)^7^ as in the previously-mentioned benchmark study^8^ (**Supplementary Fig. 12a,c**; **Methods**). Each metric measures the relative mixing of class labels, where optimal performance is associated with maximal mixing in the batch labels and minimal mixing in the cell type labels.

The “Raw” batch correction results are the first 50 principal components of the horizontally concatenated gene-by-cell expression count matrix using *stats::prcomp* in R package with centering and scaling. The “Raw, preprocessed” batch correction used the preprocessed data with log normalization with scaling factor 10^4^ and selection of 3000 highly variable genes with Seurat v3 with no restriction on the minimum number of cells and genes.

For batch correction using Seurat v3, default options are used for pancreas dataset whereas for mouse atlas dataset no cutoff was used for the minimum number of cells and genes as in Tran *et al*.^8^. The dimension of the batch corrected embedding is set as 50 dimensions following the default option for *Seurat::RunPCA* and for the consistency with SIMBA.

For batch correction using LIGER, the same arguments are used (lambda = 5, nrep = 3) are used for *liger::optimizeALS* in Tran *et al*. other than the number of factors k was set as 50 for consistency with other methods for both datasets.

For batch correction using Harmony, the same arguments are used as in Tran *et al*.^8^ other than the number of dimensions of the output embedding was set to 50 instead of 20. We note that the output embedding of 20 dimensions would result in the similar result as when used 50 dimensions in these methods.

### Supplementary Note 3: Comparison with multi-omics integration methods

Seurat3 and LIGER are two of the most widely-adopted methods for single-cell data integration. Here, we demonstrate that SIMBA outperforms these methods on two separate datasets, the recently published SHARE-seq mouse skin dataset and the similarly recent 10x PBMCs multiome dataset (**Supplementary Table 1**). We focus on Seurat3 and LIGER as they have explicit documentation for the task of integrating scRNA-seq and scATAC-seq data.

We first qualitatively evaluated these methods by inspecting UMAP visualization plots. For the SHARE-seq dataset, we observed that all three methods perform comparably well in mixing cells of two modalities though LIGER generated particularly small and noisy clusters (**Supplementary Fig. 15b)**. For the 10X PBMCs dataset, SIMBA resulted in the best mixing of cells belonging to each modality whereas other methods clustered cells separately within the originating modalities (**Supplementary Fig. 15d**). We next quantitatively assessed the integration performance of each method using four metrics that measure the distances between matched cells in the integrated space (**Methods**). In addition to the commonly-used metrics including anchoring distance, Silhouette index, and Fraction in the same cluster, we developed an additional metric, *anchoring distance rank* (ADR), which represents the normalized rank of the distance between matching cells. If two matching cells from scRNA-seq and scATAC-seq are mutually closest to one another, their ADR will be close to 0 (**Methods**) and thus a minimized ADR is ideal. Overall SIMBA showed the best performance according to ADR as well as cluster agreement while showing comparable or better performance according to the remaining metrics for both datasets (**Supplementary Fig. 15a,c**).

The modality integration procedure for Seurat v3 and LIGER follows the tutorial provided by the authors (Seurat v3: https://satijalab.org/seurat/archive/v3.1/atacseq_integration_vignette.html; LIGER: http://htmlpreview.github.io/? https://github.com/welch-lab/liger/blob/master/vignettes/Integrating_scRNA_and_scATAC_data.html).

Both Seurat v3 and LIGER formulate the modality integration task between scRNA-seq and scATAC-seq data as a batch correction task between scRNA-seq and gene activity matrix constructed from scATAC-seq. In Seurat v3, the gene activity score of a gene is calculated as the sum of the read counts in the peaks that falls within from 2kb upstream of the TSS to the end of the gene body. In LIGER, this score is calculated as the sum of all read counts that falls within 3kb upstream of the TSS to the end of the gene body.

The “Raw” results start from a scRNA-seq count matrix and a gene activity matrix calculated by Seurat v3. Filtering for the shared genes in both modalities resulted in 16738 genes for the SHARE-seq mouse skin dataset and 11045 genes for the 10X PBMCs dataset. Gene-by-cell gene expression matrix and gene activity matrix were horizontally concatenated along matching rows (genes). The output embedding is the first 20 principal components calculated by the R function *stats::prcomp* with centering and scaling.

For the modality integration using Seurat v3, the gene expression count was filtered using the default parameters *min.cells = 3* and *min.features = 200*. The co-embedding was created as described in the tutorial of the package using the scRNA-seq. The output embedding consists of the first 50 principal components, which is the default option of *Seurat::RunPCA*.

For the modality integration using LIGER, the gene expression count and gene activity matrices were normalized and filtered for the genes that are shared between both matrices. The values were then scaled according to the tutorial. In applying LIGER to the SHARE-seq mouse skin dataset, the function, *liger::optimizeALS* was used with the default parameters, k = 20 and lambda = 5. The scRNA-seq dataset was indicated as the reference in the function, *liger::quantile_norm* as described in the documentation. The scRNA-seq and scATAC-seq modalities of the 10X PBMC multiome dataset were unable to be aligned using the default parameters. Thus *lambda = 30* and *max.iters = 100* were used for the *liger::optimizeALS* function and the scATAC-seq dataset was indicated as the reference using the *liger::quantile_norm* function to ensure a better alignment.

